# A mechanistic density functional theory for ecology across scales

**DOI:** 10.1101/2021.06.22.449359

**Authors:** Martin-I. Trappe, Ryan A. Chisholm

## Abstract

Our ability to predict the properties of a system typically diminishes as the number of its interacting constituents rises. This poses major challenges for understanding natural ecosystems, and humanity’s effects on them. How do macroecological patterns emerge from the interplay between species and their environment? What is the impact on complex ecological systems of human interventions, such as extermination of large predators, deforestation, and climate change? The resolution of such questions is hampered in part by the lack of a holistic approach that unifies ecology across temporal and spatial scales. Here we use density functional theory, a computational method for many-body problems in physics, to develop a novel computational framework for ecosystem modelling. Our methods accurately fit experimental and synthetic data of interacting multi-species communities across spatial scales and can project to unseen data. Our mechanistic framework provides a promising new avenue for understanding how ecosystems operate and facilitates quantitative assessment of interventions.

---

Ecology has detailed mechanistic theories that capture the behaviour of a small number of interacting species at fine scales [1–3]. It also has broad ‘unified’ theories that ignore the details of individual species and their interactions, but can reproduce large-scale patterns, such as the relationship between number of species and area [4–7]. Despite numerous attempts, no definitive unification that bridges these theories across scales has yet been achieved. Ideally, a unified theory would be parameterisable at one scale of ecological organisation (e.g., the individual organism, or the species) and make accurate predictions at other scales (e.g., the local community or landscape). Meeting this challenge would allow knowledge from field data and limited experimental setups to be leveraged to predict the behaviour of more-complex, realistic ecosystems [4, 8–12]. Such an enterprise calls for a diverse range of quantitative approaches.

Here, we draw on density functional theory (DFT) [13–16], which has been successfully applied to a staggering number of many-body problems in physics, involving scales from atoms to molecules to planetary cores [17–20]. DFT is based on the mathematical proof that all properties of all quantum systems can, in principle, be obtained from the density distributions of its constituents. At the core of DFT lies an energy (cost) functional in terms of these density distributions. Of fundamental interest in physics are the density distributions with lowest energy (ground states and equilibria), obtained from minimising the energy over all permissible spatially varying densities. DFT predicts key properties of physical systems while automatically accounting for trade-offs between system components and constraints with efficiency and accuracy. The conceptual framework of DFT is the mathematically rigorous generalisation of the Thomas–Fermi (TF) model [21, 22] from the advent of quantum mechanics. The development and fruitful application of DFT has sky-rocketed since then, with nowadays tens of thousands of yearly publications [23]. In light of the success of DFT in simulating many-body systems within a general framework, it is worth considering whether complex systems in the life sciences can also be modelled with DFT techniques.

While there are numerous DFT applications in molecular biology [24], only one DFT-inspired study has so far emerged in macrobiology: The authors of Ref. [25] inferred spatial density distributions of *Drosophila melanogaster* (fruit flies) in an experimental arena by pairing the grand-canonical ensemble of statistical physics with an energy functional. This statistical method accurately predicts the distributions of fruit flies in new environments. Here we develop a novel and more general mechanistic approach, which we term density functional theory for ecology (DFTe). The core of DFTe is an energy functional for ecosystems that merges ecological principles with insights from DFT, especially from the TF model of multi-component quantum gases. Our DFTe bridges ecological scales in an intuitively accessible, comprehensive framework and successfully predicts ecological data from systems with a variety of interacting taxa and environmental constraints.

## RESULTS

Many key questions in ecology, such as abundance redistribution in response to environmental change, ask for constrained equilibria — a goal shared by DFT in physics. At the core of DFT lies a cost function, the density functional of the energy *E*. Once the building blocks of *E* are specified, its minimisation delivers the position-(***r***-)dependent equilibrium (viz., ground-state) densities ***n***(***r***) = {*n*_1_(***r***)*, n*_2_(***r***), …, *n_S_* (***r***)} for the abundances ***N*** = {*N*_1_*, N*_2_, …, *N_S_*} of any interacting many-body system of *S* species, such as a gas of several atomic species.

We have transferred the DFT narrative and methodology to ecology by building a quantum-gas-inspired mechanistic energy functional

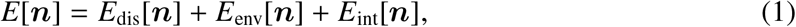

whose minimisation balances species-specific costs due to dispersal, environment, and interactions, including resource consumption (see Methods). In referring to *E* as an energy for a given realisation (***n, N***) of an ecological system, we quantify a relation to other configurations (***n′, N′***). The insight that a system aims at occupying the state of lowest (ground-state) energy among all possible system configurations is one of the fundamental principles of physics. The ecological (axiomatic) equivalent comes in various forms, with the *statistical* maximum entropy theory of ecology being one of them [26]. Here, we construct the *mechanistic* DFTe energy *E* such that (i) its global minimum realises the ecosystem equilibrium with abundances 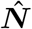 and (ii) its geometry away from 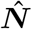 gives rise to both steady-state and nonequilibrium dynamics.

The energy of a system is a measure of its ability to invoke change in another system. Twice the energy, relative to the other system, doubles this ability — energy is additive. But this does not imply that the energy of a system decomposes into a sum as in equation (1). We base the additive decomposition of *E* on our understanding of how constituents of physical systems, in particular of interacting quantum gases, come together to assemble the energy. For the dispersal energy *E*_dis_ we adopt the TF kinetic energy expression (see Methods) — ultimately based on the quantum-mechanical spin-statistics theorem, which prohibits two identical fermions from occupying the same state [27] and which therefore implies an intra-specific pressure that implements one form of dispersal. But this declaration of *E* does not introduce any quantum effects into ecology. We rather determine the functional form of *E* by introducing and building on intuitive concepts *in analogy* to the energy components of physical systems. Furthermore, the structurally identical DFT counterpart of *E* in physics is rigorously founded in quantum-mechanical principles, but, in ecology, we do not pursue the hopeless goal of establishing DFTe based on actual energetic quantities and mechanisms. Exploiting the mathematical structure of and tools developed for DFT, we rather formulate DFTe as a computational framework by establishing an energy functional that is informed by ecological principles and benchmarked not against energies but against data of densities and abundances.

Among the various implementations of DFT, we find that density-potential functional theory (DPFT) [28–33] is uniquely qualified for achieving our objectives. Combined with the TF model, DPFT delivers the equilibrium density

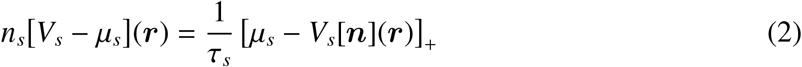

of species *s*, with species-specific dispersal pressure constants *τ_s_* (which tend to dilute *n_s_*, see Methods and Extended Data Fig. 1). The Lagrange multipliers ***μ*** in equation (2) are used to enforce ∫_*A*_(d***r***) ***n***(***r***) = ***N*** in a focal area *A* (see Methods). The operator []_+_ stems from the TF model and ensures nonnegative densities by replacing negative arguments with zero. The potential energy

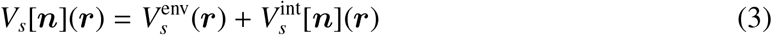

effectively merges the environment 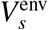 (as perceived by species *s*) with the interaction potential 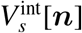 (the functional derivative of *E*_int_[***n***]), which couples *s* to the other species. We omit arguments of functions for brevity wherever the command of clarity permits.

The coupled equations (2) and (3) are the essence of DPFT in general and of DFTe in particular: According to equation (2), the members of species *s* seek to reside where the cost *V_s_* due to environment and interactions is small, while the self-consistent solution of equations (2) and (3) produces an automatic trade-off against intra-specific dispersal pressure (see Methods). Until further evidence suggests otherwise, we deem the TF-based equation (2) sufficient for our phenomenological DFTe, which leaves us with creating suitable interaction functionals *E*_int_[***n***] for ecology (see Methods).

In the following, we enlist five biological systems to build and verify the computational DFTe framework. We thereby (i) demonstrate the alignment of DFTe results with both empirical and synthetic data, (ii) discuss advantages as well as limitations of DFTe relative to existing modelling approaches in ecology, (iii) illuminate how to deploy the building blocks of DFTe in practice, and (iv) show how to extract problem-adjusted DFTe tools from the general DFTe energy for studying modifications and extensions of ecosystems to which our theory has been fit. Figure 1 summarises our agenda by listing the ingredients and main insights of the DFTe predictions for our five systems. We find that the performance of DFTe is similar to that of existing system-targeted modelling approaches, see Figs. 2, 3, and 5. Beyond this, a major benefit of DFTe is its generality.

**FIG. 1.**
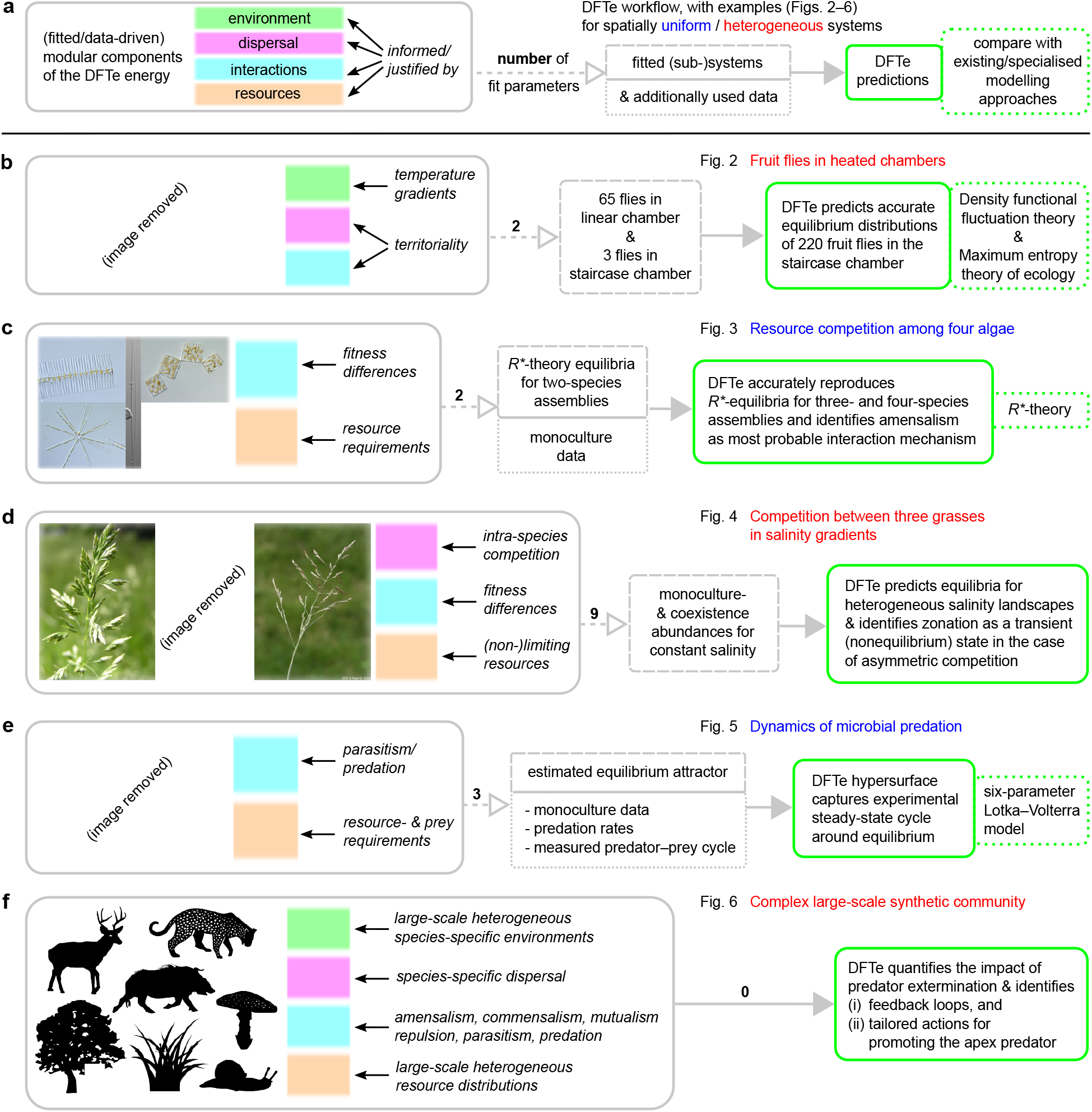
Agenda for establishing, validating, and applying DFTe. **a**, The workflow for extracting DFTe predictions from ecosystems whose properties justify and inform our choices for the components of the DFTe energy functional *E* (see Methods). Fitting the resulting explicit parameterisations of *E* to less complex subsystems, we create problem-adjusted tools for addressing the complete systems and, more importantly, their modifications and extensions. **b**–**f**, We highlight our main results for five experimental and synthetic systems (Figs. 2–6), which cover a considerable range of taxa, interactions, and environmental settings. The successful modelling of these diverse setups gives hope that our DFTe framework can address a much broader range of ecological systems, and that it is the first stepping stone towards a universal density functional theory for ecology as a whole. In view of the conceptual disparity between DFTe and existing ecological theories, it is expedient to make connections at the level of predictions rather than the basic equations: We argue that DFTe adds generality to ecosystem modelling, but is usually not expected to outperform specialised approaches. Details on the parameterisation of each example are spelled out in Results and Methods. See Supplementary Information for details on image sources.

**FIG. 2.**
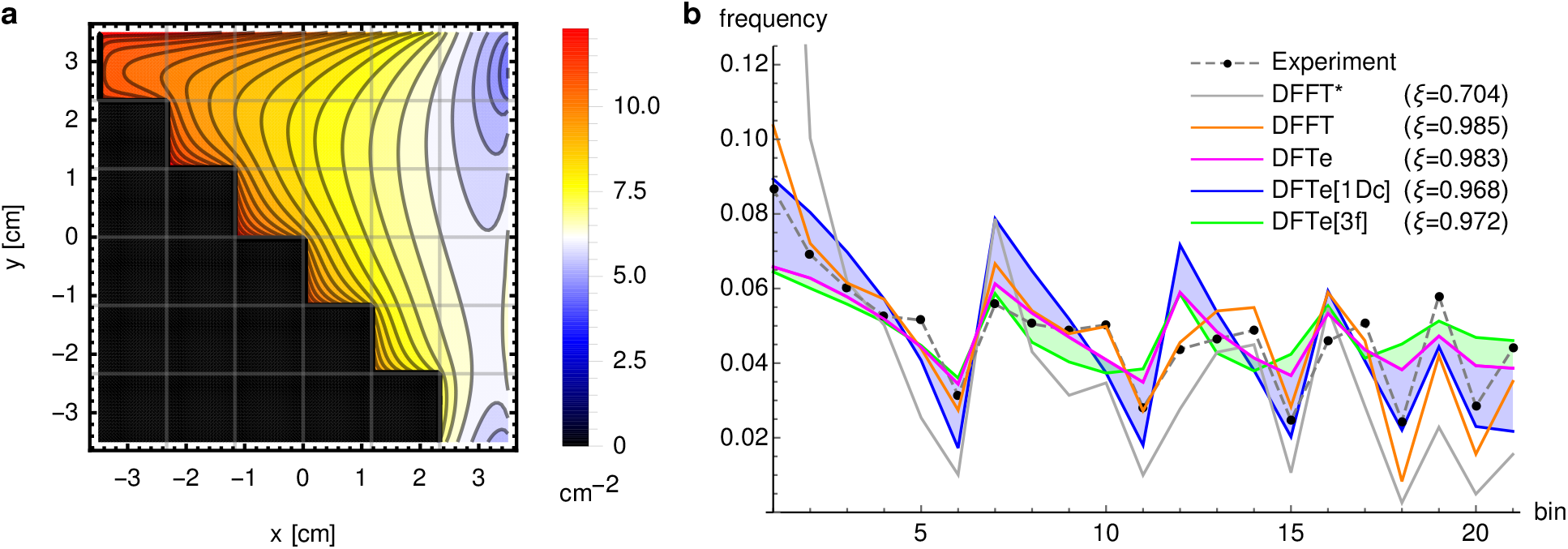
An application to confined fruit flies demonstrates how DFTe predicts spatial distributions of fixed abundances. **a**, The DFTe prediction of the spatial density distribution of 220 interacting flies in a staircase chamber with heat source at *x* = 3.5 cm. **b**, We benchmarked against the experimental data by extracting the average of the continuous DFTe density profile in each of the 21 grid cells accessible to flies (in row-major order starting from the top left bin of **a**). Measuring the overlap of predictions and data via the least-squares correlator *ξ*, we found that DFTe achieves essentially the same accuracy (*ξ* ≈ 1) against the experimental data as DFFT [25] — using the same data for fitting.

**FIG. 3.**
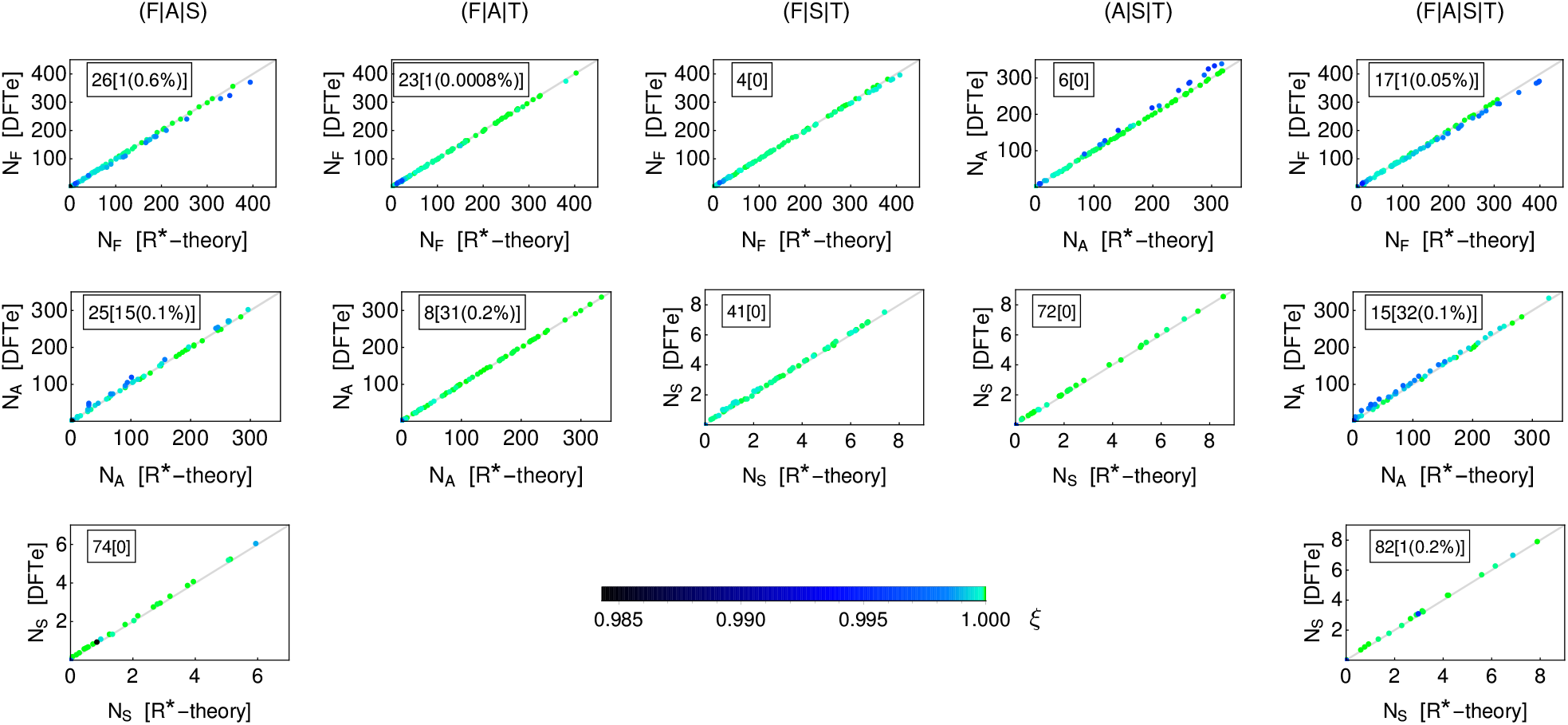
An application to Tilman’s algae competition experiments demonstrates DFTe’s ability to handle resource constraints. Without exception, we found that DFTe delivers the *R*^∗^-equilibria of the four-species assemblage (F|A|S|T) and its three-species subsets accurately (*ξ* ≈ 1; the colour legend applies to all graphs). Each of the five columns of graphs represents 100 randomly drawn resource combinations and shows the resulting DFTe abundances *N_s_* (*s* ∈ {F, A, S, T}; given per micro-litre of suspension) for one of the five algae assemblages relative to *R*^∗^-predictions. The DFTe abundances are represented by coloured points, each associated with one resource combination. If both DFTe and *R*^∗^-theory predict the same abundance of one of the system’s species, then the coloured point falls on the diagonal gray line of the corresponding graph, while its colour reports the overlap *ξ* of all (three or four) abundances with the *R*^∗^-predictions for the associated resource combination. We do not show results for T, which is competitively excluded in all 500 cases within both *R*^∗^-theory and DFTe. The framed box within each graph displays the number of resource cases (out of 100) for which the species was excluded in (i) both *R*^∗^-theory and DFTe, and (ii) in *R*^∗^-theory but not in DFTe [in square brackets; the percentage gives the (on-average) abundance of the minority species relative to the majority species, that is, the minority species was almost excluded]. We observed the latter case only if both F and A, which have similar competitive abilities (see Supplementary Information), were part of the assemblage. Supplementary Fig. 2 shows a summarising histogram in terms of *ξ*.

The fundamental computational procedure of DFTe is the minimisation of the energy functional *E* for fixed abundances ***N***. We demonstrated this with our first example, the fruit fly system of Méndez-Valderrama et al. [25], where our DFTe is able to predict distributions of flies in novel situations with similar accuracy as state-of-the-art methods (Fig. 2). This example illustrates the interplay between dispersal, environment, and interactions in DFTe.

Following the DFTe workflow in Fig. 1b, we fitted a two-parameter DFTe energy functional *E* (equation (24) in Methods) to data of 65 flies in a crowded heterogeneous environment, a heated elongated chamber (‘1Dc’-data), see Extended Data Fig. 1, and to an uncrowded heterogeneous environment with three flies in a staircase chamber exposed to a heat source (‘3f’-data). We then used *E* to predict the equilibrium distribution of 220 fruit flies in the same staircase chamber (Fig. 2a) and thereby determined the trade-offs between dispersal, environment, and interactions (see also Extended Data Fig. 1). We normalised this density distribution to one and assessed the deviation between (i) the resulting frequency values (labelled ‘DFTe’ in Fig. 2b) for the 21 bins of the staircase chamber and (ii) the experimental frequencies. For that purpose we used the least-squares correlation measure *ξ* = 2 ***p*** · ***d***/(***p***^2^ + ***d***^2^), which quantifies the overlap of the vectorised predictions ***p*** (e.g., from DFTe) with the data ***d*** (here, the experimental data). Figure 2b shows that DFTe yields similar accuracies (*ξ* ≈ 1) as density-functional fluctuation theory (DFFT), which is developed in Ref. [25] as a statistical approach based on the experimental data shown here and, unlike DFTe, cannot identify mechanisms of the kind implied by equation (24). We also report two alternative DFTe predictions obtained from fitting exclusively to the 1Dc- and 3f-data, respectively. These are labelled ‘DFTe[1Dc]’ and ‘DFTe[3f]’ in Fig. 2b and reproduce the experimental data less accurately than the ‘DFTe’ outcome, which incorporates both the 1Dc- and the 3f-data. However, all three DFTe predictions lie within the blue- and green-shaded areas, which illustrate the spread of potential DFTe outcomes if we were oblivious to the optimal data-informed fitting procedure (see Methods) or if only one of the two data sets were available. That is, all three DFTe predictions perform significantly better than the ‘naive’ DFFT^∗^ model, which essentially scales the density distribution of three flies up to 220 flies. The latter scaling behaviour would also result from the maximum entropy theory of ecology if solely constrained to the 3f-data: Without system-specific information beyond the spatial three-flies distribution, a maximum entropy evaluation has to assume independent individuals and thus reproduces the DFFT^∗^ outcome.

Next, we demonstrated how DFTe can incorporate the constraints of resource availability on species’ coexistence. In seminal resource competition experiments, D. Tilman showed that the outcomes can be predicted by fitting a dynamic model (*R*^∗^-theory) to data from single-species experiments [1]. In Fig. 3, we showed that DFTe accurately reproduces Tilman’s predictions of the equilibrium abundances of four algae in an experimental lab setting by supplying the DFTe energy components (equation (28) in Methods) with species-specific resource requirements and fitness proxies based on the monoculture data of Ref. [1]. This illuminates the trade-off between resources and interactions in DFTe. In this case study, we benchmarked DFTe against the equilibrium abundances that follow from *R*^∗^-theory (see Methods and Extended Data Tab. 2), rather than against the (transient) experimental data of Ref. [1]. With a single fit of the two free parameters of the DFTe energy functional to all the outcomes of resource competition according to *R*^∗^-theory [1] among all two-species subsets of the four algae *Fragilaria crotonensis* (F), *Asterionella formosa* (A), *Synedra filiformis* (S), and *Tabellaria flocculosa* (T) for a comprehensive set of resource combinations, we built a tool for describing more complex situations, such as all three- and four-species assemblages for any resource supply. We report the according results in Fig. 3. Instead of incorporating all single-species parameters of the microscopic *R*^∗^-theory, and thus following the difficult path of highly parameterised microscopic descriptions for multi-species systems [34], we used this example to illustrate the principal usage of DFTe as an effective description, whose crucial input are outcomes of measurements on simple systems. Specifically, we parameterised the DFTe energy solely by arguing that the algae’s competitive abilities depend on the inverses of (i) the minimum resource requirements (*R*^∗^-values) and (ii) the nutrient requirements per cell (see Methods). This application of DFTe required fitting of two free parameters at the two-species level, while *R*^∗^-theory is completely parameterised at the single-species level. This highlights one limitation of DFTe that comes with the higher level of abstraction. We also dis-regarded the parameters of resource flow rates ***F*** and maximal growth rates in *R*^∗^-theory: While equilibria for different ***F*** are readily available through *R*^∗^-theory, they cannot be obtained by the DFTe energy fitted to data for fixed ***F***.

In all our simulations, the *R*^∗^-predictions for *S* > 2 were consistent with a recently studied assembly rule that predicts coexistence in microbial multi-species communities from the survivors of two-species competition [35]. Figure 3 shows that DFTe predicts the same patterns of competitive exclusion, though in many cases, the inferior species is merely strongly suppressed instead of excluded, which highlights DFTe’s capacity of extracting system properties accurately (but not necessarily exactly) from a continuum of a priori possible outcomes.

Next, with an application to three grasses in a saline environment [36], we showed how DFTe can project sparse information from uniform experimental setups to density distributions in heterogeneous environments (see Methods). One non-trivial result from our DFTe simulation is zonation, i.e., the occurrence of sharp boundaries between species distributions despite a continuous environment [37]. In Fig. 4 we identify zonation as a transient, interaction-induced phenomenon, which may even persist as an itinerant steady state if environmental disturbances prohibit complete equilibration. These predictions can potentially be tested in field studies.

**FIG. 4.**
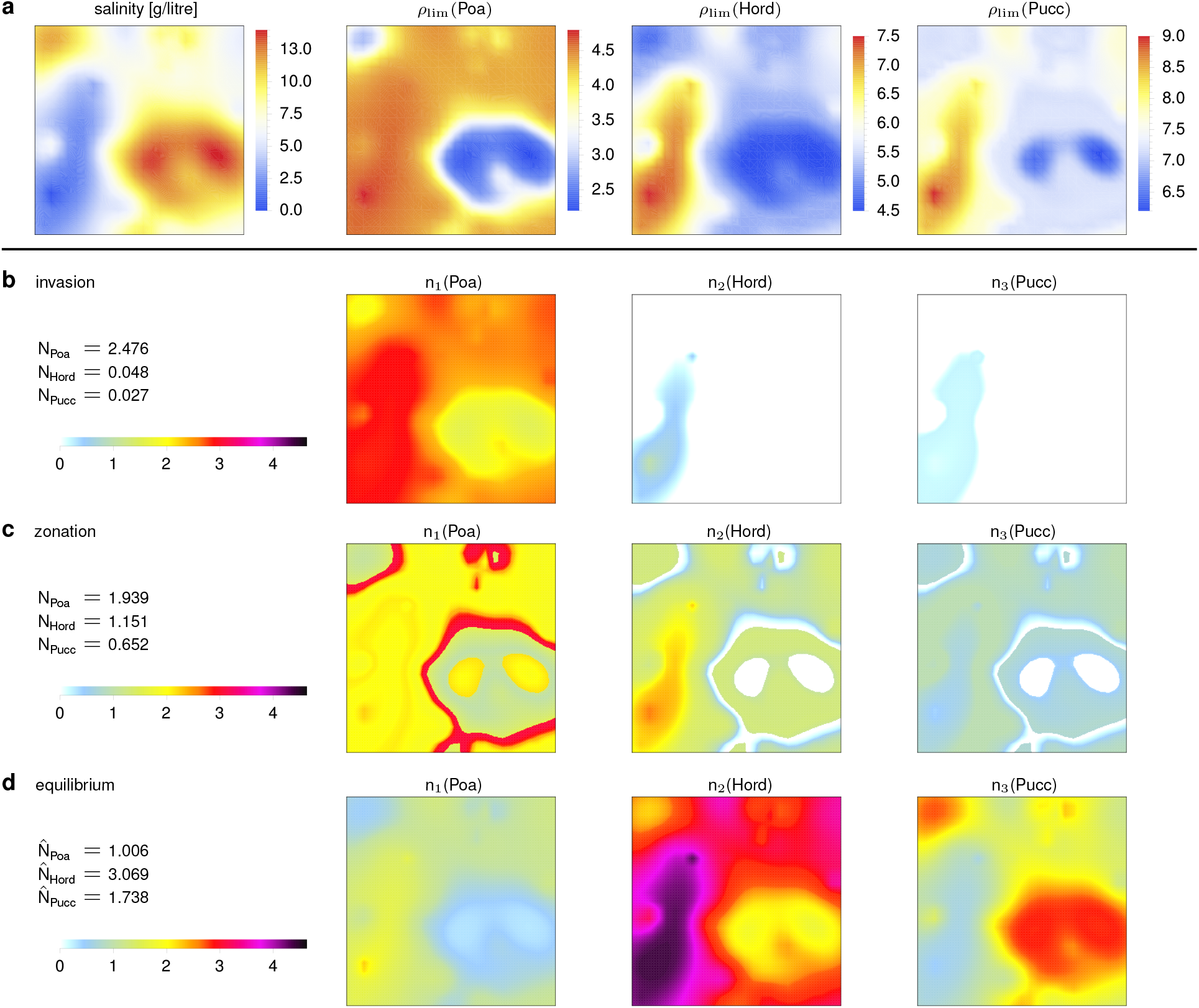
An application to plants in salinity grad1ients demonstrates how DFTe extrapolates scarce experimental information into realistic settings. **a**, We fitted the DFTe functional (equation (29) in Methods) with asymmetric repulsive contact interactions between the three grass species *Poa pratensis* (Poa), *Hordeum jubatum* (Hord), and *Puccinellia nuttalliana* (Pucc) to above-ground biomass data. These data are reported in Ref. [36] for monoculture and mixture setups at spatially uniform salinity levels and translate a randomly generated landscape with heterogeneous salinity (square area *A* = 1) into distributions of the hypothetical, but data-informed, limiting resources *ρ*_lim_ for each species. **b**, A subsequent DFTe simulation showed how Hord and Pucc invade Poa’s equilibrium monoculture distribution. We predicted a rich zoo of phases (see Methods and Extended Data Fig. 2), including zonation as a transient state (**c**) on the way from Poa’s initial monoculture distribution (Extended Data Fig. 2a) to the smooth equilibrated mixture (**d**).

For our first two systems, the fruit flies and the competing algae, we focussed on comparing the DFTe equilibria to data. But for our third system, the competing plants in a saline environment, DFTe also generated non-trivial predictions of transient dynamics (Fig. 4). With a fourth example, we now demonstrate that DFTe can also model nonequilibrium steady states, such as predator–prey dynamics. We thereby showed that the DFTe hypersurface 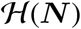, which is the space of DFTe equilibria spanned by all permissible ***N*** (see equation (14) in Methods) and is specified through system-specific instances of equation (1), has predictive power also away from the equilibrium abundances 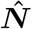, although 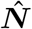 may a priori follow from a myriad of energy functionals *E*[***n***, ***μ***](***N***) that share the same global minimum.

Figure 5 illustrates our DFTe results on the microbial predator–prey system of *Didinium nasutum* feeding on *Paramecium aurelia*, where experiments have indicated that the long-term steady state is not an equilibrium but cycles [38, 39]. We found that the observed predator–prey cycles are accurately captured by the DFTe energy functional (equation (30) in Methods). This opens the door to modelling time-dependent phenomena in ecology with DFTe (see Supplementary Notes).

**FIG. 5.**
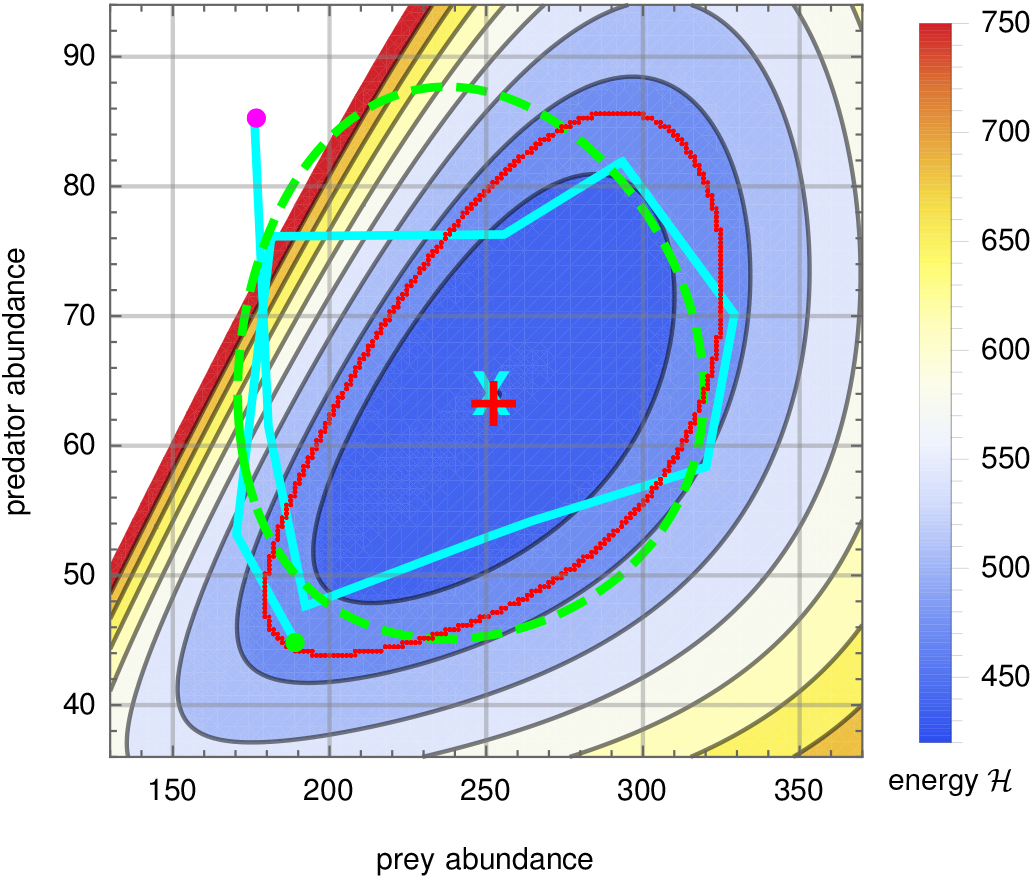
An application to a predator–prey system demonstrates the ability of DFTe to capture steady-state dynamics. The DFTe hypersurface 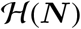 is a platform for ecosystem dynamics, much like the analogous energy surface of classical physics is for Kepler orbits in a gravitational field. The cyclic DFTe equipotential line (red curve) probes 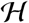 away from equilibrium (red plus sign) and captures the empirically measured cycle (cyan curve, starting at the magenta dot and ending at the green dot) centred on the cyan cross, which represents the average measured abundances and anchors the DFTe fit. Strong excitations on 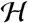 permit predator extinction along noncyclic trajectories: The high-energy equipotential lines terminate at the abscissa, but not at the ordinate (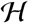 exceeds 750 in the white region and diverges towards the ordinate; see also Extended Data Fig. 3b for an overview plot of 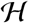). The DFTe trajectory lacks information on the time-resolved rates of abundance changes and, thus, does not translate unambiguously into a time series that would allow point-wise comparison with predictions from dynamic models — see Supplementary Information for ways to augment DFTe with explicit Newtonian-Type time evolution based on 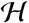. However, we compute the DFTe trajectory by fitting only three parameters and find it to be of similar quality to a fitted trajectory from the standard six-parameter Lotka–Volterra model (dashed green; see Supplementary Information), as judged by the total area between the experimental cycle and each of the predicted trajectories.

Finally, we explored the full potential of DFTe with the aid of a synthetic community of seven interacting species (Fungus, Deer, Pig, Snail, Tree, Grass, Cat; Fig. 6) subjected to heterogeneous environments and resource distributions that are reminiscent of, for example, regional-scale distributions. The density *n_s_* of each species may, independently from the other densities *n*_*s′ ≠ s*_, refer to individuals per area, frequency, biomass density, fractional land cover, or any other expedient metric — units and absolute scales are absorbed in the parameters *τ_s_*, 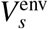, and so forth. By deploying only contact-type interactions (see Extended Data Tab. 3), we implicitly state that members of any species are not directly influenced by conspecifics or heterospecifics beyond their local pixel of our coarse-grained area *A*. For example, if the Cat habitat area is ~ 50 km^2^, then *A* exceeds ~ 50, 000 km^2^ (see Supplementary Notes). This large-scale system contrasts with the fruit flies experiment in Fig. 2, where the long-range repulsive interaction between individuals turns out to couple all individuals explicitly across the entire area. Note that contact-type interactions on large spatial scales do not imply that the system is weakly interacting: Both the synthetic food web here and the fruit flies experiment are strongly interacting since the interaction energies comprise a substantial part of the total energy for both systems.

**FIG. 6.**
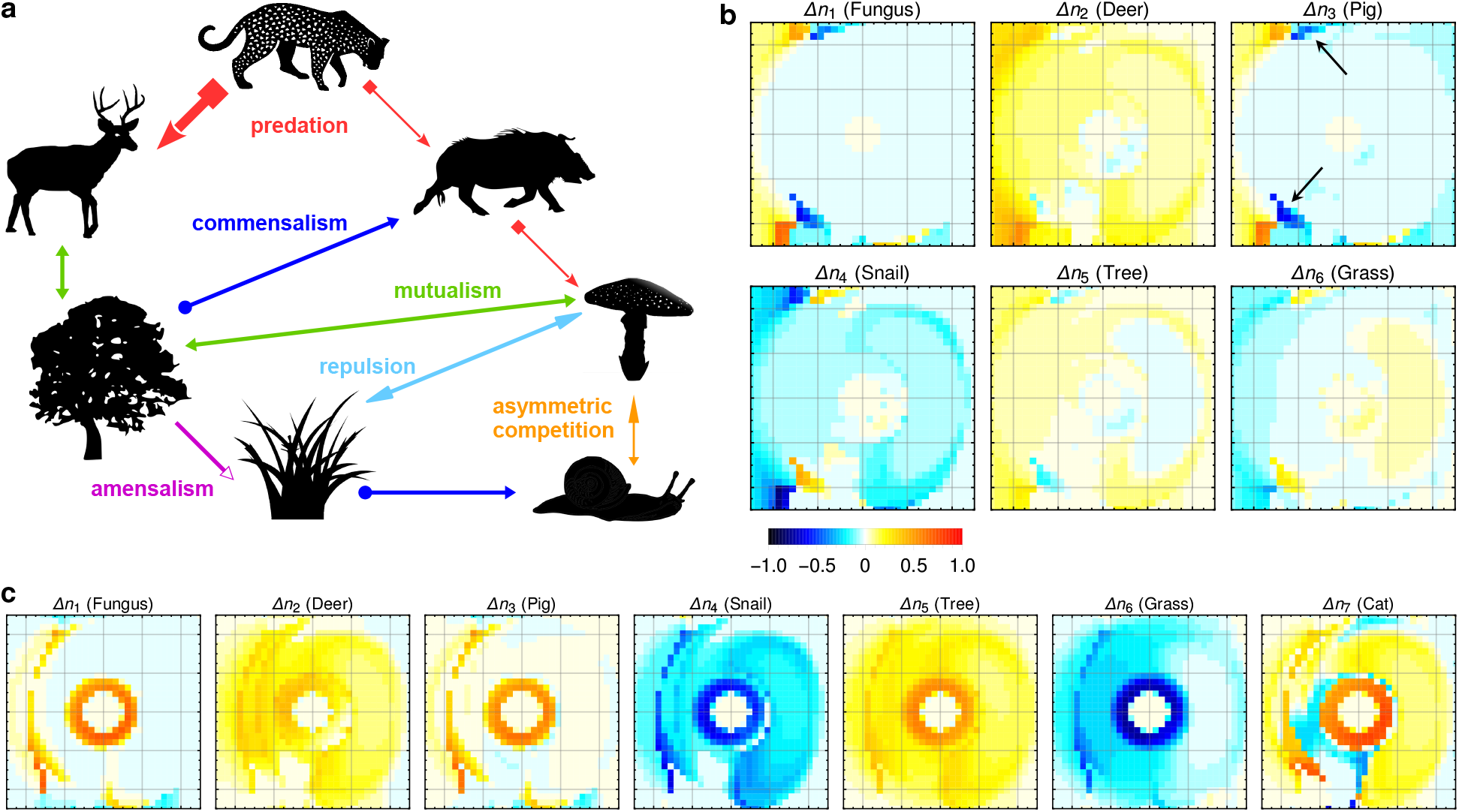
An application to a hypothetical food web demonstrates DFTe’s abilities to make predictions about the effects of perturbations on complex ecosystems. **a**, The cartoon illustrates the food web involving seven hypothetical species. **b**, The six sub-panels depict the relative changes of species distributions following the extermination of the Cat (see graphical legend; redder colours indicate increasing populations; bluer colours indicate decreasing populations). While density surges of the Cat’s prey (the Cat requires both Deer and Pig) are to be expected, the absence of the apex predator has repercussions through-out the entire community — evidently, the interaction-mediated links in the community are strong enough to induce major distortions of the density distributions of all species. This includes effects that may come as a surprise prior to our quantitative simulation, such as regionally declining Pig populations. Two such enclaves are indicated with arrows. We found the main effect of removing the Cat to be a re-equilibration of the whole ecosystem through feedback loops that promote Fungus, Deer, and Tree, but penalise Grass and Snail. We gained confidence in these DFTe predictions by successively building the complete community from simpler subsystems that permit an intuitive understanding of the mechanistic relations between the model ingredients and the equilibrated density distributions (see Methods and Extended Data Fig. 6 and Supplementary Notes). For a real system, we could then use the quantitative knowledge obtained from the perturbation simulation to promote the Cat through informed actions: **c**, Cutting the Tree’s deforestation stress in half, we increased the Cat population by 36% and created new Cat habitat, especially in the central ring region (see Methods and Extended Data Fig. 7).

Using DFTe, we quantified the impact of removing the apex predator and thereby highlighted the capacity of DFTe for predicting the effects of perturbations on complex multi-species communities in realistic spatially structured environments [40]. Despite the complexity of this multi-trophic system, we retain ample intuition of the causal relations between the model input and output by successively building the complete ecosystem. This understanding extends to effects that may appear counterintuitive when examining only the interactions in Fig. 6a. For example, in the following we explain the small decrease of the Pig abundance upon Cat extermination. First, we observed that the Pig is the limiting resource for the Cat only in regions of very low Pig density (see Extended Data Fig. 6), such that changes in the Pig distribution are not primarily connected to the removal of the Cat. Second, the heavily predated Deer as well as the mutualistically connected Tree generally benefit from the absence of the Cat. As a result, the Grass and, consequently, the Snail come under pressure, which helps the Fungus, whose mutualistic connection to the Tree closes the positive feedback loop between Deer, Tree, and Fungus. The increased Fungus density then attracts the Pig, except in two small enclaves (indicated with arrows in Fig. 6b), where the processes just described are reversed, such that the Pig is mainly redistributed globally. Of course, this narrative conveys only the broad strokes of the quantitative simulation and plays out under the constraints imposed by environments and resources, but it conveys the quantitative knowledge about the community functioning obtained from the DFTe simulation. In a practical setting, this would allow us to promote the apex predator through informed and tailored actions, here for example by reducing the Tree’s deforestation stress (see Extended Data Fig. 4). This demonstrates the potential of DFTe for informing management strategies for biodiversity, for which purpose food web models are currently commonly consulted. Our modular and spatially explicit DFTe framework overcomes major challenges of traditional food web analyses, which often lack (i) the crucial incorporation of relevant processes beyond inter-species interactions and (ii) spatial heterogeneity [41], with some exceptions that have been emerging recently (for example, Ref. [42]) or that are designed for specific ecosystems (for example, Ref. [43]).

## DISCUSSION

Ecology has a long tradition of modelling simple systems with mechanistic equations, for example, of the Lotka–Volterra type. Some properties of large and complex ecosystems have also become accessible through statistical approaches. Here we have presented a single mechanistic framework that facilitates bridging across scales — a goal that has been recognised as both elusive and important for addressing fundamental ecological questions. Inspired by techniques from density functional theory (DFT) in physics and models of quantum gases, we have created a density functional theory for ecology (DFTe), an entirely novel approach to theoretical ecology, and demonstrated its alignment with experimental and synthetic data across various ecological systems. We have not yet found a case study of which DFTe does not provide an excellent description, although we encourage further work exploring more case studies.

The key advantage of DFTe over existing approaches to ecosystem modelling lies in its capacity to incorporate all relevant processes on all spatial scales, while the complexity of the underlying DFTe energy functional, which mirrors that of the target ecosystem, is of little technical concern for retrieving the density distributions and the trade-offs between the involved processes. Unlike, for example, the maximum entropy theory of ecology [26] and the density-functional fluctuation theory [25], which also root in equilibrium principles of physics, DFTe is a mechanistic theory. DFTe features an intuitive and straightforward conceptual foundation, ease of technical implementation, and scope of applicability, unifying the major ecological principles [44] in a single mechanistic framework. DFTe also contrasts with the established mechanistic theories of ecology, which usually neglect essential processes and whose rigid equations become ever harder to solve and to revise with rising system complexity [3]. Analogous to DFT applications in physics, we have shown that the components of an ecosystem’s DFTe functional can be inferred independently from simpler subsystems. This resolves the ubiquitous problem of finding the proper trade-offs among given mechanisms and constraints for many species that interact within a complex environment. The ecological constituents and mechanisms are independently parameterised components of, and coupled through, the DFTe functional, whose composite structure permits hassle-free revision and introduction of causal mechanisms that can correct disparities between predictions and data. For example, higher-order interactions [45, 46] can simply be added to equation (9) in Methods. Densities and abundances may refer to individuals or aggregated observables like biomass or land cover. Conversely, we may fork one species into many, for example, when accounting for heterogeneity among individuals is expedient [9, 47].

As within any mechanistic framework, and in contrast to some statistical theories like the maximum entropy theory of ecology, the predictive power of DFTe is limited when simulating poorly understood systems or if critical data are missing. Important practical challenges towards a universally useful DFTe are thus (i) the selection of the most important components of the energy functional, which is expedient for including as few fitted parameters as possible, and (ii) the required basic intuition on how to combine the ecologically relevant parameters into an energy functional expression. However, with the examples of Figs. 2, 3, and 5, we have highlighted that rudimentary insight into ecosystem functioning can be sufficient for generating results on a par with established ecological theories. Judging from the work presented here, we deem spatial resolution to be the only technical challenge in scaling up DFTe to simulate, for instance, hundreds to thousands of species on planet Earth at 1 km^2^-resolution, which likely requires high-performance computing at dedicated facilities — assuming that enough data for such a scenario is available to parameterise the DFTe energy sufficiently.

Having successfully modelled diverse systems from predatory microbes in test tubes to contact-interacting grasses in saline landscapes to a synthetic food web, we have demonstrated the potential of DFTe for application to a broad range of ecological problems. While we are optimistic that DFTe can become a universal tool for an intuitive approach to ecology as a whole, including dynamics on all relevant time scales (see Supplementary Notes), only in-depth future analyses can be the ultimate judge on that matter.

On a general note, we hope that our foundational work at the cross-roads of physics and ecology inspires researchers and practitioners across disciplines towards developing DFT-based tools that tackle their big questions, such as pandemics, traffic control, or predicting the dynamics of financial assets. These are serious enterprises, and a concerted effort will be required. We hope to start an interdisciplinary dialogue that transfers one of the most powerful methods of physics to unexpected realms.

## METHODS

### Modular components of the DFTe energy functional

The central ingredient of DFTe is an energy functional *E*, assembled according to equation (1). Specifically,

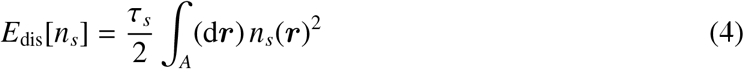

is the two-dimensional analogue of the TF kinetic energy functional for fermionic quantum systems and encodes intra-species pressure of species *s*, e.g., individuals repel conspecifics from territory; note that energy density represents pressure in general. *E*_dis_ implies a conspecific negative density dependence, relative to other energy components whose integrands scale less than quadratically with density. We emphasise that we neither aim at describing all forms of dispersal via equation (4) nor necessarily need to include *E*_dis_ even if a concrete ecosystem exhibits dispersal, which can potentially be modelled with other energy components. Akin to the external energy in physics, the environmental energy

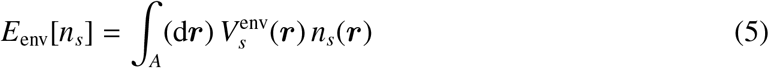

parallels the parameterisation in Ref. [25] and models, for example, habitat preference and external influences, such as spatially varying climate and local deforestation stress. Figure 2 demonstrates that equation (5) can realistically describe the influence of the environment on species’ distributions. Back reactions on the environment can be introduced as part of *E*_int_, through a bifunctional *E*_br_[***V***^env^, ***n***] that yields the equilibrated modified environment 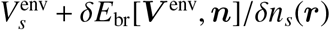, cf. equation (3). Summing equations (4) and (5) over *s*, we obtain the respective total energy components displayed in equation (1). Inspired by analogues of interacting quantum gases, we let *E*_int_[***n***] include all possible bipartite interactions

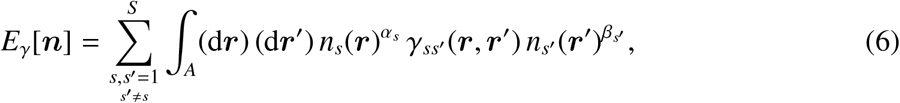

which include amensalism, commensalism, mutualism, and so forth. Here, *α_s_, β_s′_* ≥ 0, and the interaction kernels *γ_ss′_* are assembled from fitness proxies of species *s* and *s*′ (Extended Data Tab. 1). Higher-order interactions can be introduced, for example, through (i) terms like *n_s_ γ_ss′_ n_s′_ γ′_s′ s′′_ n_s′′_* that build on pairwise interactions, cf. Ref. [45], or (ii) genuinely multipartite expressions like *γ_ss′ s′′_ n_s_ n_s′_ n_s′′_*. For (*α_s_, β_s′_*) = (1, 1) we identify the contact interaction in physics as *γ_ss′_* ∝ *δ*(***r*** − ***r′***) with the two-dimensional delta function *δ*(), while the Coulomb interaction amounts to setting *γ_ss_* ∝ 1/|***r*** − ***r***′|. The mechanistic effect of these interaction kernels on the density distributions is the same in ecology as it is in physics, and both feature in describing the ecosystems addressed in this work. While the contact interaction is a suitable candidate for plants and especially microbes [48], we expect long-range interactions (for example, repulsion of Coulomb type) to be more appropriate for species with long-range sensors, such as eyes.

In a natural setting the equilibrium abundances are ultimately constrained by the accessible resources. It is within these limits of resource availability that environment as well as intra- and inter-specific interactions can shape the density distributions. An energy term for penalising over- and underconsumption of resources is thus of central importance. Each species consumes resources from some of the *K* provided resources, indexed by *k*. A subset of species consumes the locally available resource density *ρ_k_*(***r***) according to the resource requirements *ν_ks_*, which represent the absolute amount of resource *k* consumed by one individual (or aggregated constituent) of species *s*. The simple quadratic functional

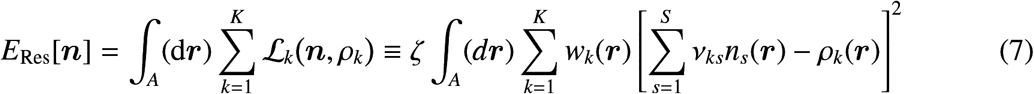

proves appropriate. Here, *ν_ks_n_s_* is the portion of resource density *ρ_k_* that is consumed by species *s*. That is, *ν_ks_* > 0 indicates that *s* requires resource *k*. Predator–prey relationships are introduced by making species *k* a resource *ρ_k_* =]*n_k_*[, where]*n*[declares *n* a constant w.r.t. the functional differentiation of *E*, that is, the predator tends to align with the prey, not the prey with the predator. In view of the energy minimisation, the quadratic term in equation (7) entails that regions of low resource density *ρ_k_* are less important than regions of high *ρ_k_*. The different resources *k* have the same ability to limit the abundances, such that the limiting resource *k* = *l* at ***r*** has to come with the largest of weights *w_l_*(***r***), irrespective of the absolute amounts of resources at ***r***. For example, the weights *w_k_* have to ensure that an essential but scarce mineral has (a priori) the same ability to limit the abundances as a resource like water, which might be abundant in absolute terms. To that end, we specify the weights

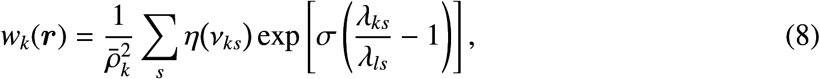

which are inspired by the smooth minimum function, where *σ* < 0, *λ_ks_*(***r***) = *ρ_k_*(***r***)*/ν_ks_*, and the carrying capacity of the limiting resource is 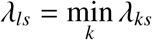. The step function *η*() in equation (8) ensures that only resources *k* that are actually consumed by species *s* contribute to *w_k_*. We rescale *w_k_* using the average 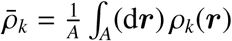, which puts all resources on equal footing in their ability to limit abundances and renders *E*_Res_ invariant under change of resource units. The ratio of carrying capacities in equation (8) implements the relative importance of all resources at ***r***, with resources *k* ≠ *l* suppressed exponentially according to their deviation from their ability to limit the abundances. For example, with *σ* = −4 and *λ_ks_* = 2*λ_ls_*, resource *k* ≠ *l* is largely irrelevant for species *s* as it weighs in at less than 2% compared to the limiting resource *l*. There can be multiple limiting resources for a species *s*, most conceivably, multiple resources that vanish locally. In the limit of vanishing resource density (*ρ_l_, λ_ls_* → 0), the exponential in equation (8) reduces to the Kronecker-delta *δ_kl_*, thereby rendering all resources with *λ_ks_* > *λ_ls_* irrelevant at ***r***. Using *E*_Res_, we show that an analytically solvable minimal example of two amensalistically interacting species already exhibits a plethora of resource-dependent equilibrium states (see Supplementary Notes and Supplementary Fig. 1).

We specify the DFTe energy functional in equation (1) by summing equations (4)–(7) and (optionally) constraining the abundances to ***N*** via Lagrange multipliers ***μ***:

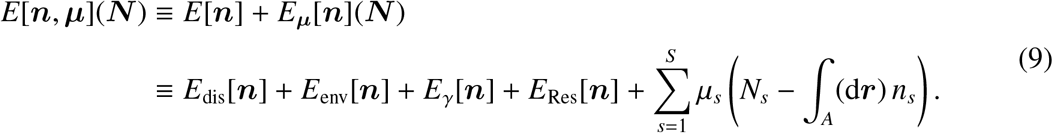

Uniform situations are characterised by spatially constant ingredients *n_s_* = *N_s_/A*, *ρ_k_* = *R_k_/A*, coefficients *τ_s_*, etc. for the DFTe energy, such that equation (9) reduces to a function *E*(***N***) with building blocks

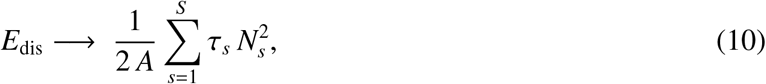

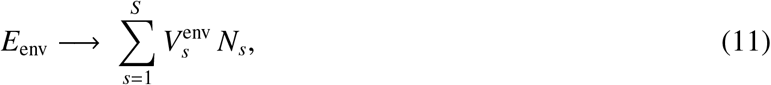

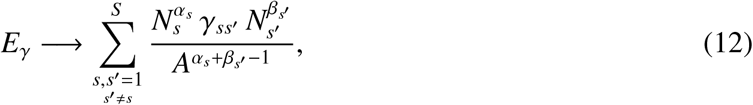

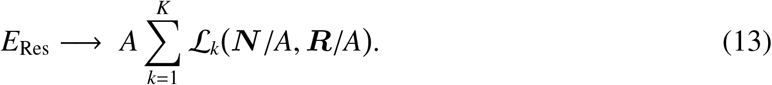

### Ecosystem equilibria from the DFTe energy functional

The general form of equation (9) gives rise to two types of minimisers (viz., equilibria): First, we term

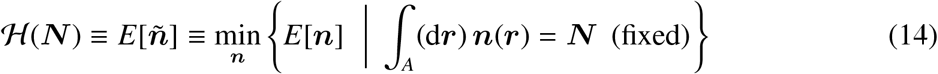

the ‘DFTe hypersurface’, with 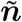 the energy-minimising spatial density profiles for given (fixed) ***N***. Second, the ecosystem equilibrium is attained at the equilibrium abundances 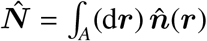 which yield the global energy minimum

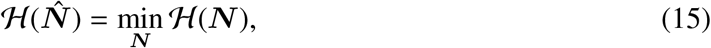

where the minimisation samples all admissible abundances, that is, 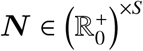 if no further constraints are imposed.

The direct minimisation of *E*[***n***] is most practical for uniform systems, which only require us to minimise *E*(***N***) over an *S* -dimensional space of abundances. For the general nonuniform case, we adopt a two-step strategy that reflects equations (14) and (15). First, we obtain the equilibrated density distributions on 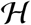 for fixed ***N*** from the computational DPFT framework [28–33]. Second, a conjugate gradient descent searches 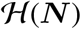 for the global minimiser 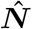. Technically, we perform the computationally more efficient descent in ***μ***-space. Local minima are frequently encountered, and we identify the best candidate for the global minimum from many individual runs that are initialised with random ***μ***. Note that system realisations with energies close to the global minimum, especially local minima, are likely observable in reality, assuming that the system can equilibrate at all.

### Density-potential functional theory (DPFT) in Thomas–Fermi (TF) approximation

Defining

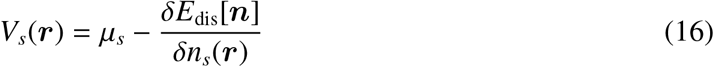

for all *s*, we obtain the reversible Legendre transform

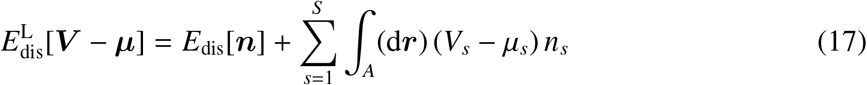

of the dispersal energy and thereby supplement the total energy with the additional variables ***V***:

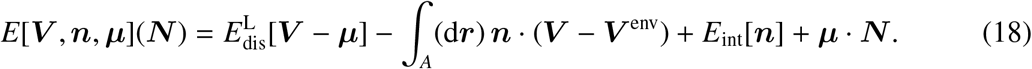

This density-potential functional is equivalent to (but more flexible than) the density-only functional *E*[***n***, ***μ***](***N***). The minimisers of *E*[***n***] are thus among the stationary points of equation (18) and are obtained by solving

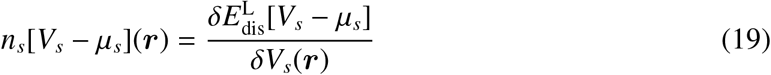

and

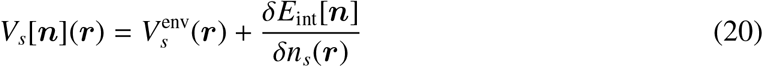

selfconsistently for all *n_s_* while enforcing ∫_*A*_(d***r***) *n_s_*(***r***) = *N_s_*. Specifically, starting from ***V***^(0)^ = ***V***^env^, such that 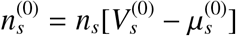, we iterate

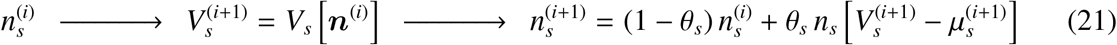

until all *n_s_* are converged sufficiently. This selfconsistent loop establishes a trade-off between dispersal energy and effective environment ***V*** by forcing an initial out-of-equilibrium density distribution to equilibrate at fixed ***N***. We adjust 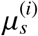 in each iteration *i* such that 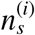 integrates to *N_s_*. Small enough density admixtures, with 0 < *θ_s_* < 1, are required for convergence. Each point of 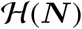 in equation (14) represents the so equilibrated system for given ***N***.

Assuming a given interaction energy, we require an explicit expression for the right-hand side of equation (19) to realise the selfconsistent loop of equation (21). For two-dimensional fermion gases in TF approximation [30], we have

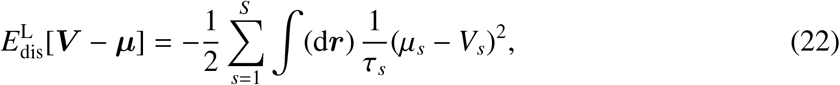

which delivers (i) the density in equation (2) to be used for the right-hand side of equation (19) and (ii) equation (4) via equation (17) upon inverting the functional relationship *n*[*V*] of equation (2). For stabilising the numerics if necessary or if, for example, unambiguous derivatives of the density are sought, we replace equation (2) by its smooth version

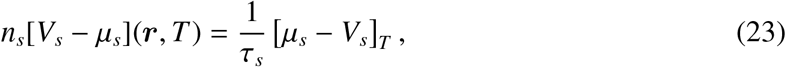

where [*x*]_*T*_ = *T* log 1 + exp *x/T* is a smooth version of [*x*]_+_. If *τ_s_* = 0, we can add a dispersal energy with positive *τ_s_* to *E* in order to execute the selfconsistent loop with equations (2) or (23), and compensate by subtracting the same dispersal energy from the interaction energy, which enters equation (20).

### Explicit parameterisations of the DFTe energy functional

*Figure 2.* In view of equation (2), an educated guess informed by the measured reference density 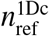 (see Extended Data Fig. 1) is a quadratic environment induced by the heat source at *x*_0_. We also include a repulsive long-range interaction, since (i) the data in Fig. 2g of Ref. [25] suggests that the interaction is quadratic in the density and (ii) fruit flies have to sense their peers remotely, for example, in defending territory [49]. This leaves us with two parameters, *ε* and *γ*, for fitting the minimising density of

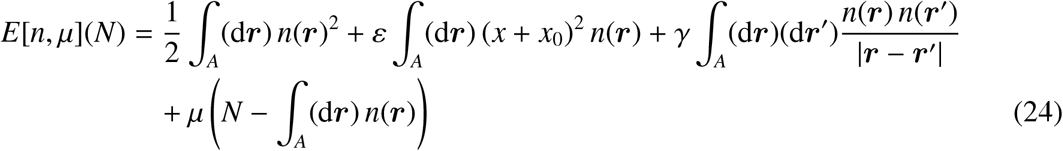

to 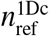. In Ref. [25] the interaction parameters from the quasi-one-dimensional (1D) chamber are combined with the environmental information from a low-density experiment (three flies) in a ‘staircase chamber’ to predict a high-density distribution of 220 flies (labelled ‘DFFT’ in Fig. 2b). This is a reasonable procedure, since the low-density three-flies experiment represents a situation with very small total repulsion and is therefore well suited for extracting the environmental influence on the density distribution. We thus follow the same strategy. Keeping *γ* = 9 cm from the density fit to the quasi-1D chamber setup, we get *ε* = 14.5 cm^−2^ for three flies in the staircase chamber and use both parameters for predicting the density distribution of 220 flies (labelled ‘DFTe’ in Fig. 2b). Since *γ* is (i) one of the system’s characteristic length scales, (ii) an estimator of the spatial reach of the long-range interaction, and (iii) close to the system size, we deem the fruit flies setup a small-scale system — in contrast to the large-scale contact-interacting systems of Figs. 4 and 6.

*Figure 3.* We face a uniform environment with *S* species and two resources. An amensalistic interaction is an appropriate candidate for putting a subset of species at a disadvantage relative to their heterospecifics (see Extended Data Tab. 1). We rule out parasitism, understood in the sense of an asymmetric interaction, for which the DFTe equilibria do not produce the correct survivors in all reference resource cases 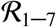 (see Extended Data Tab. 2). In contrast to the fruit-flies study above, the abundances are not fixed but are rather to be determined as minimisers of the ***N*** -dependent energy function *E*(***N***) = *E*_Res_(***N***) + *E_γ_*(***N***), assembled from equations (10)–(13). The spatially uniform interaction kernel

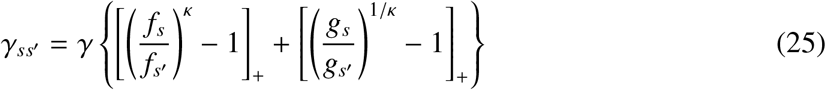

for *E_γ_*(***N***), see equation (12), is assembled from the fitness proxies

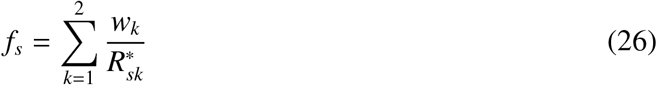

and

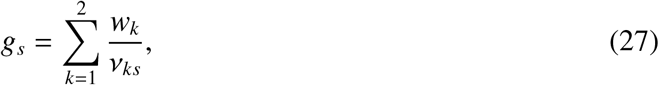

which we assume to influence the species’ ability to consume the provided resources in the presence of heterospecifics. The *s*-specific traits 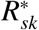 (the densities of resource *k*, below which species *s* cannot survive in monoculture) and *ν_ks_* see equation (7) follow from monoculture experiments (see Supplementary Notes and Tab. 1 of Ref. [1]). The exponent *κ* in equation (25) introduces a hierarchy between the fitness proxies of equations (26) and (27) and serves as a second parameter to be used in the fitting of our reference data, which are all 42 ‘reference’ abundances 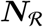 of two-species *R*^∗^-equilibria that follow from 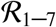. The parameter *γ* enables the trade-off between competition and resource energy. In the spirit of *R*^∗^-theory, we choose *σ* → −∞ in equation (8), such that only the limiting resource for each species enters the weights *w_k_* for *E*_Res_. Finally, the minimisers of

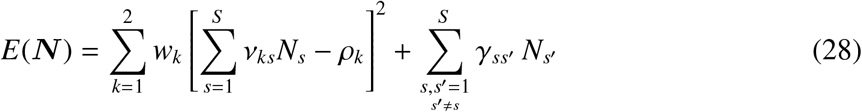

yield the best fit to 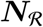 for *γ* = 8 × 10^−8^ and *κ* = 8, used for modelling the three- and four-species communities in Fig. 3.

*Figure 4.* The slim characterisation of the experimental setting reported in Ref. [36] (see Supplementary Notes) contrasts with the number of parameters of the DFTe functional *E*[***n***, ***μ***](***N***) in equation (9), for which all components require input — except the environmental potential energies, which are constant in this uniform situation and therefore irrelevant. In order to proceed, we assume the following: (i) Coexistence of the three competing grasses in mixture suggests that three different resources are exclusively limiting each species; (ii) all species exhibit equal dispersal *τ* as well as interaction strength *γ* factored into *γ_ss′_* = *γ* [*f_s_/ f_s′_* − 1]_+_ — mixture- and monoculture abundances are not proportional, implying some kind of salinity-dependent competition; (iii) asymmetric interactions, since amensalism and repulsion yield distributions inconsistent with field data, see Supplementary Notes and Supplementary Fig. 3; (iv) as suggested in Ref. [36], we fix the salinity-dependent fitness proxies *f_s_* as the fraction of above-ground biomass of *s* in mixture. This reflects increasing salt tolerance in the order Poa < Hord < Pucc. We then fit the nine parameters (*τ,σ,ν*_*k*≠*s*_, *γ*) of

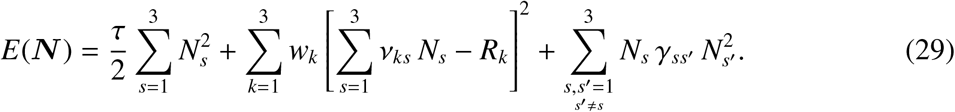

to the abundances for the mixture in uniform salinity (see Supplementary Figs. 3 and 4), such that the nonuniform version of equation (29) allows us to predict the density distributions in Fig. 4 implied by heterogeneous resource distributions (see Supplementary Notes).

*Figure 5. Paramecium aurelia* (P) feeds on Cerophyl (c) and serves as the single resource for the predator *Didinium nasutum* (D). From the general DFTe energy we thus select the two quadratic resource energies and a parasitic interaction with fit parameter *γ*, akin to an asymmetric repulsive contact interaction that favours D. For this spatially uniform microbial system in suspension, *E*[***n***] reduces to the function

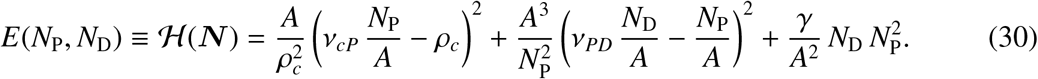

We fit its minimiser 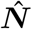 (red cross in Fig. 5) to the average abundances (cyan cross in Fig. 5) for the last cycle of the most stable time series (Fig. 14c in Ref. [38]); see also Supplementary Notes, Supplementary Fig. 5, and Supplementary Tab. 1. These average abundances anchor our comparison between the experimental data and the DFTe trajectory 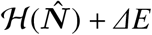, with a suitably chosen ‘excitation’ energy *ΔE*. As expected, amensalistic interactions are not supported by the data and are therefore ruled out, see Extended Data Fig. 3 and Supplementary Fig. 6.

*Figure 6.* The community encompasses heterogeneous environments, competition over resources, all bipartite interactions of Extended Data Tab. 1, and predator–prey relations. From equation (9), we assemble the according DFTe energy functional

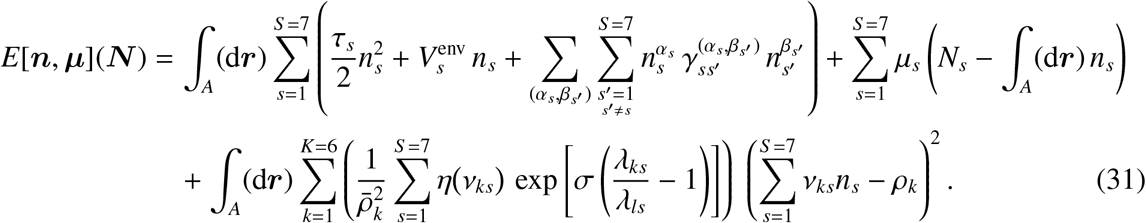

The species differ in their dispersal strengths *τ_s_*, habitat preferences 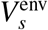, the types (*α_s_, β_s′_*) of heterospecific interactions, modulated by interaction strengths 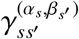, as well as the types and amounts of required resources, see Extended Data Tab. 3. The synthetic environments and non-prey resource distributions are depicted in Extended Data Fig. 4. Using the so specified energy functional along with the density expression of equation (2), we simulate 33 × 33 = 1089 parcels in the focal area *A* = 1. We obtain the equilibrium density profiles from a conjugate gradient descent towards the global energy minimum of equation (31) in the up to seven-dimensional space of abundances ***N***. In view of the stark disparity between species distributions (i) in isolation and under the influence of interactions, see Extended Data Fig. 7, we build intuition on the entire community by modelling subsystems with fewer species (Extended Data Figs. 5 and 6).

## DATA AVAILABILITY

The experimental data used in this manuscript are extracted from Refs. [1, 25, 36, 50]. The fruit-flies data from Ref. [25] are available at https://doi.org/10.1038/s41467-018-05750-z. We digitised Fig. 6 of Ref. [1], available at https://doi.org/10.2307/1937747, for displaying the experimental data on the competing algae in Extended Data Tab. 2. For our case study on three grasses, we obtained the above-ground biomass data in monoculture and mixture by digitising Fig. 3 of Ref. [36], available at https://doi.org/10.1139/b91-310. For our predator–prey case study we used the digitised data of Ref. [38], which is provided in the appendix of Ref. [50], available at https://doi.org/10.1098/rspb.2000.1186.

## CODE AVAILABILITY

All the codes used in this study are available as open access repositories. The DFTe formalism for the heterogeneous systems is part of the C++ software package ‘mpDPFT’, available at https://doi.org/10.5281/zenodo.4774448. Analyses of the uniform systems are available as Mathematica notebooks at https://doi.org/10.5281/zenodo.4774456.

## ACKNOWLEDGMENTS

We are grateful to James Patrick O’Dwyer for many insightful discussions and his crucial review of this work. We also thank Berthold-Georg Englert, Daniel Heyen, and Rosita Mei Ling Koh for valuable feedback on this manuscript. This research was supported by the ISF-NRF Singapore joint research program (grant number WBS R-154-000-B09-281).

## AUTHOR CONTRIBUTIONS

Both authors M.-I. T. and R. A. C. conceived the study and wrote the manuscript. M.-I. T. designed and carried out the research and analyses. R. A. C. conducted the Lotka–Volterra simulation and contributed to the choice of case studies for the research as well as to the interpretation of the analyses.

## EXTENDED DATA

**Extended Data Tab. 1.**
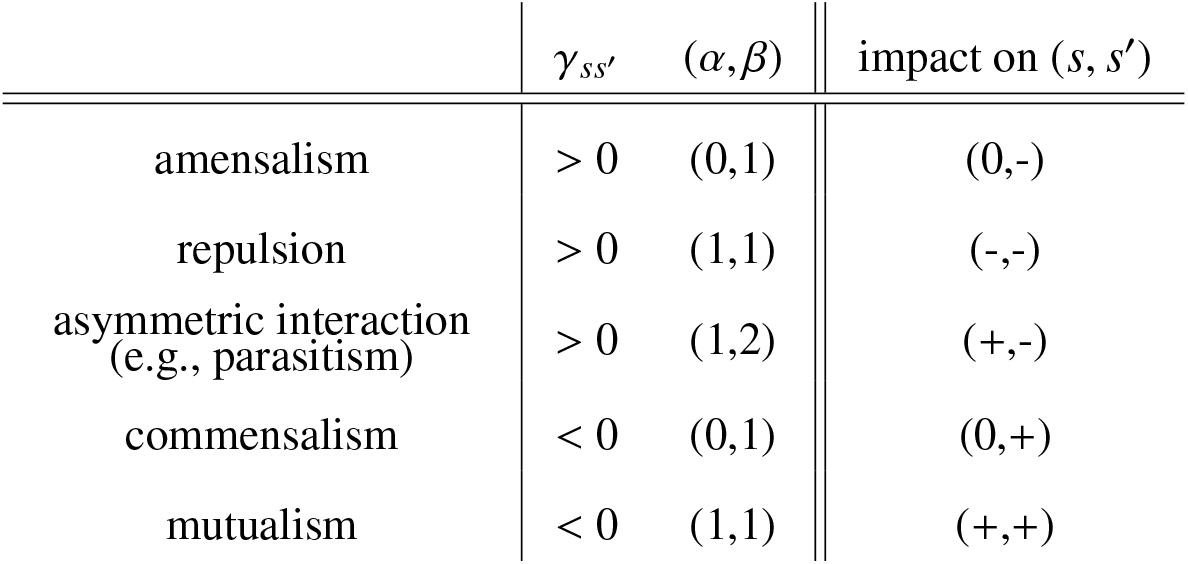
Bipartite interactions. All basic interactions in ecology are covered with the generalised competition energy *E_γ_* in equation (6), which favours (+), penalises (−), or leaves the species unaffected (0). A competitive advantage of one species *s* over another species *s*′ requires lower contribution to the energy if *n_s_* > *n_s′_*. Indeed, for *γ_ss′_* = *γ_s′s_* > 0 and *β* > *α*, we find *s* favoured over *s*′ if 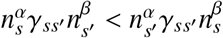, which is equivalent to 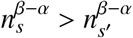. That is, the asymmetric competition incurs a cost for the superior species, but a larger cost for the inferior species. For *α* = *β*, the interaction is symmetric in *s* and *s*′, such that an overlap of *n_s_* and *n_s′_* is energetically favoured (punished) for *γ_ss′_* < 0 (*γ_ss′_* > 0). For amensalism and parasitism, we may, for example, specify a generic coefficient such as *γ_ss′_* = [*f_s_/ f_s′_* − 1]_+_ ≥ 0, with species-specific fitness proxies ***f***.

**Extended Data Fig. 1.**
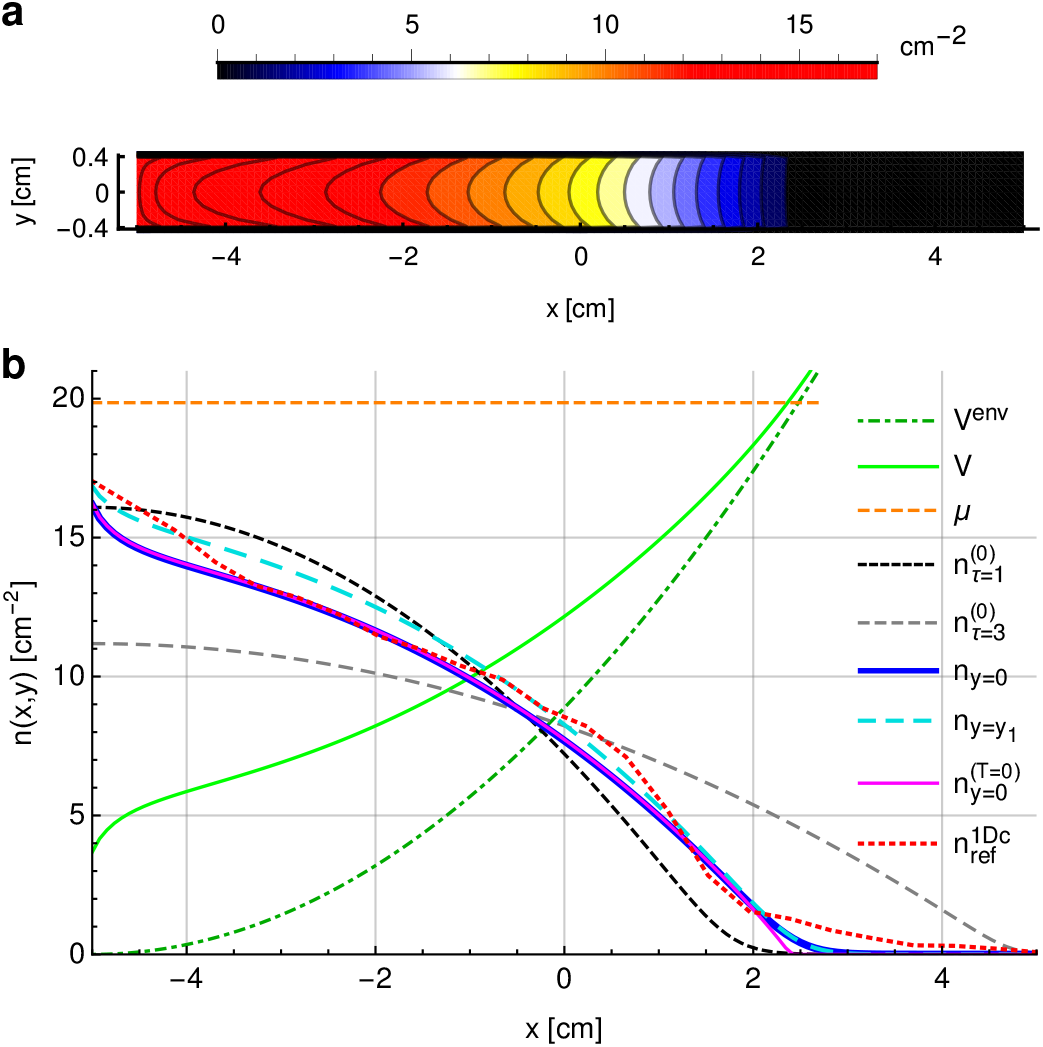
Interpretation of the TF density from DFTe fits to fruit flies. **a**, Equation (24) with *ε* = 35.5 cm^−2^ and *γ* = 9 cm yields the DFTe equilibrium densities of 65 fruit flies in the quasi-1D chamber (‘1Dc’; area *A* = 10 × 0.8 cm^2^; heat source at *x* = 5 cm; from Ref. [25]) as least-squares fits to the experimental data interpolated by 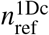. A priori, *τ*, which enters equation (24) via equation (4) is a fit parameter as well, but a rescaling of the energy does not affect its minimiser. We thus may set 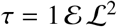, in our units of energy [*ε* = 1] and length [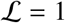 dm = 10 cm], since we only seek densities and relative energy differences. **b**, The energies *V*, *V*^env^, and *μ* are shown rescaled by 1/100. With the quadratic environment in equation (24), the TF density formula in equation (2) produces an inverted parabola for the density distribution prior to interactions — a first approximation of 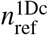. The cuts *n_y_* along the x-direction for *y* = 0 and *y*_1_ = 0.391 cm give an average account of 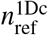. Owing to the nonlocal repulsion, the density increases towards the chamber boundaries at *y* = ±0.4 cm, but this is a small effect and of no qualitative importance here: The density profiles along the x-direction are a reasonable fit to 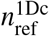 for all *y*. We obtain the results in Fig. 2 using equation (23) with *T* = 100, which produces a smooth version of 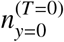 (viz., equation (2)) in the vicinity of the habitable zone boundary (the analogue of the quantum-classical boundary) at *V* = *μ*. In the absence of interactions in equation (24), *V* equals *V*^env^ and yields 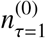 (for *ε* = 35.5 cm^−2^), which reveals the repulsive nature of both the interaction and the dispersal energy when compared with *n_y_*_=0_ and 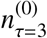, respectively.

**Extended Data Tab. 2.**
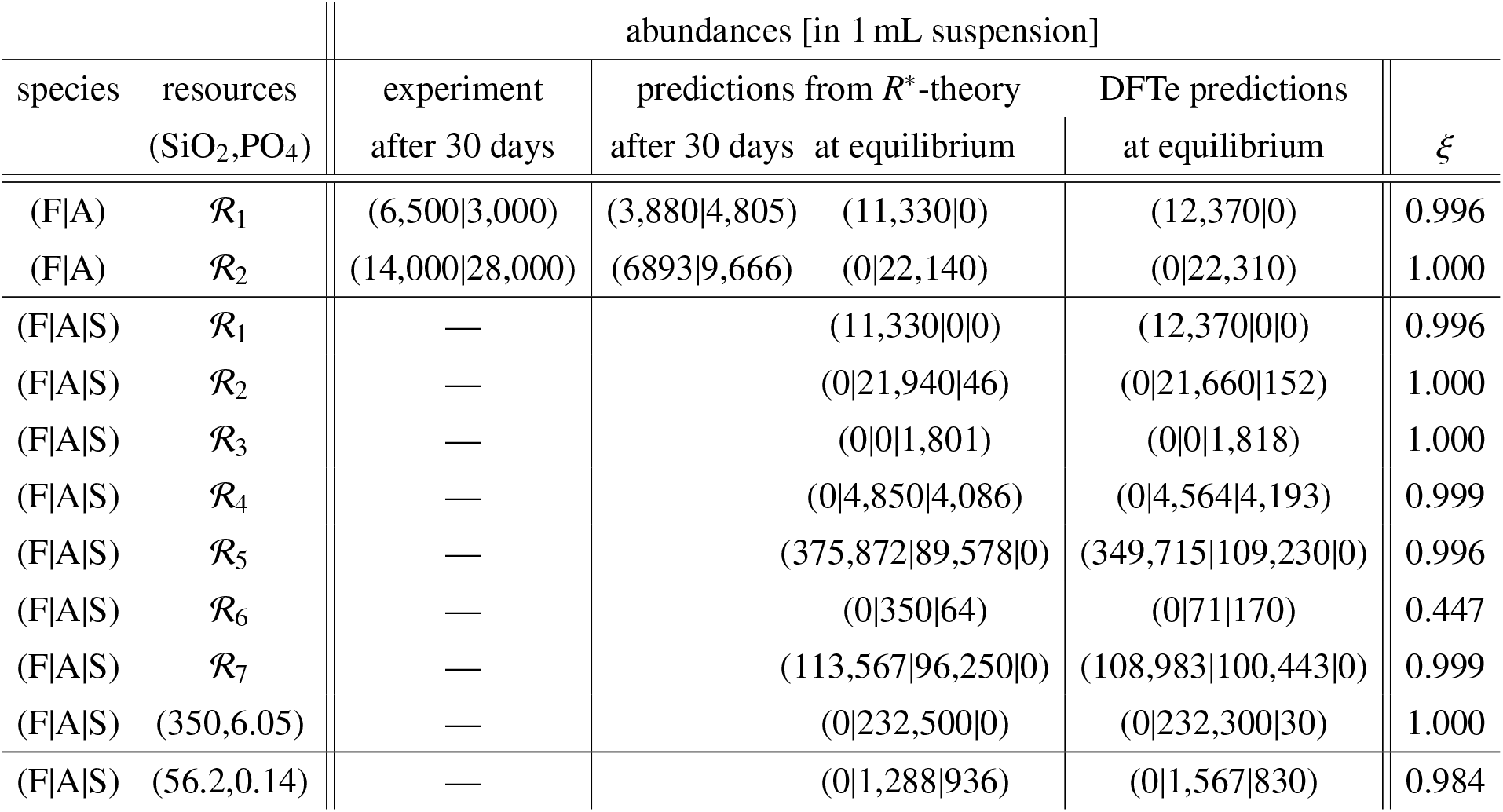
Analysis of *R*^∗^-predictions for benchmarking DFTe. The (F|A) subset of the four species (F|A|S|T) studied in Ref. [1] suggests to benchmark the DFTe trade-off between resources and competition against *R*^∗^-equilibria instead of the nonequilibrated experimental abundances after 30 days. Furthermore, the resource combinations 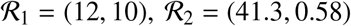, and 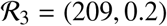 in units of *μ*mol/L, used for the experiments in Ref. [1] do not lead to strong signatures of coexistence: Competitive exclusion prevails apart from the two borderline cases 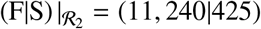 and 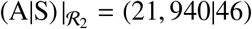, where S is almost excluded by F and A, respectively. We cover a broader range of abundances by considering four additional resource combinations (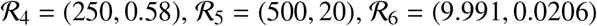, and the average 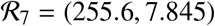 of 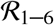) that result in indisputable coexistence according to *R*^∗^-theory. In all our calculations, DFTe never predicts coexistence of three species, in line with *R*^∗^-theory. Since T is always excluded, the most interesting assemblage is (F|A|S), then equivalent to (F|A|S|T) at equilibrium, where each species is able to competitively exclude the others depending on the provided resources. DFTe also shows that A can coexist with either F or S, while F and S cannot coexist — in agreement with *R*^∗^-theory. Our results match the *R*^∗^-predictions quantitatively. The bottom row for (F|A|S) details the case of smallest overlap *ξ* between DFTe- and *R*^∗^-equilibria found among all 500 of our randomly chosen resource combinations, see Fig. 3. The observed discrepancies between DFTe and *R*^∗^-theory are most visible when the resource combinations are very close to the *R*^∗^-values of the involved algae (e.g., 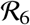 for (F|A|S)). These disparities are accounted for by the fact that *E*_Res_ permits complete resource consumption (see Supplementary Notes).

**Extended Data Fig. 2.**
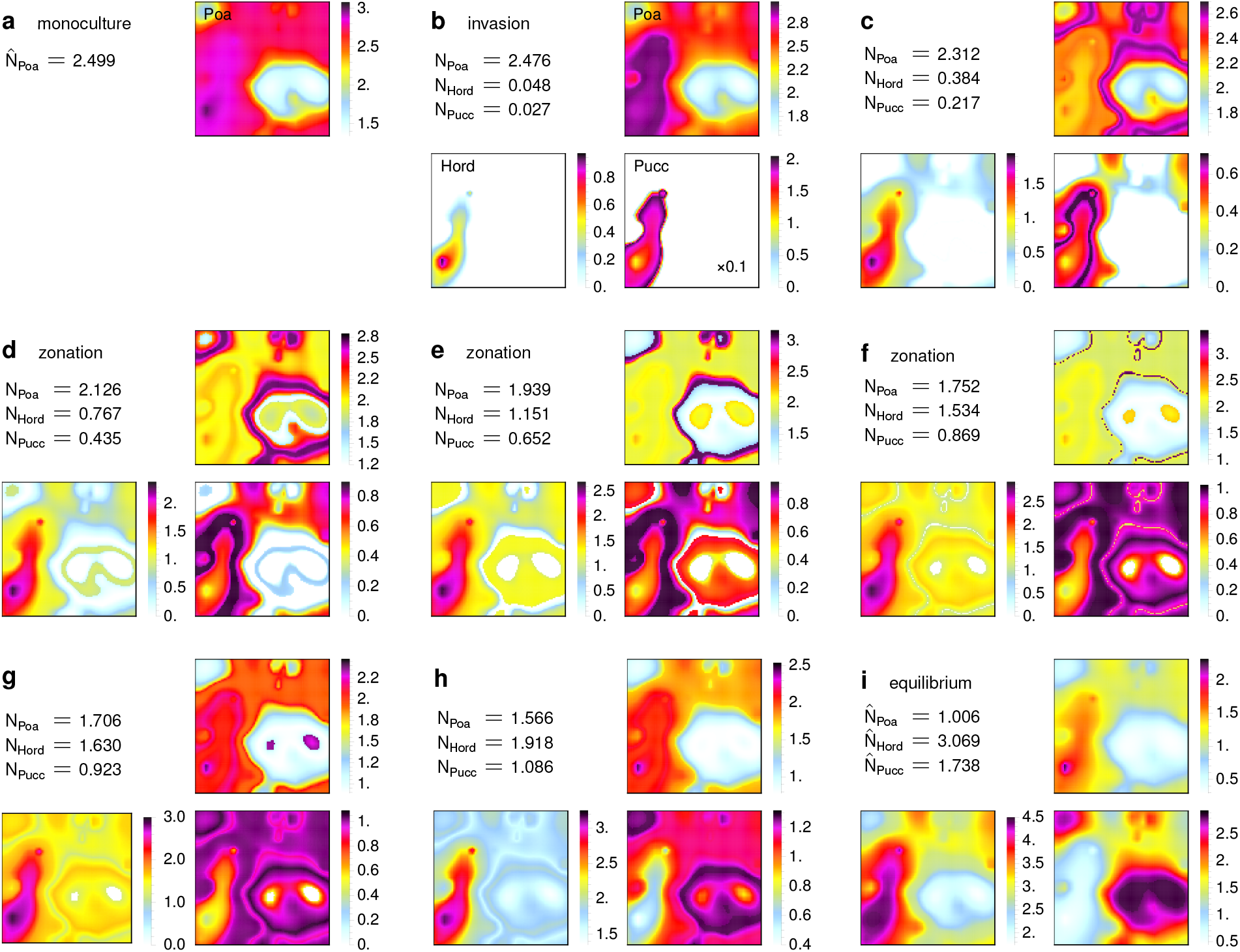
Three grasses on a trajectory to equilibrium. The density distributions of Poa, Hord, and Pucc exhibit a zoo of phases that emerge, transform, and disappear as we trace out a straight line in ***N*** -space, from Poa’s monoculture distribution (**a**) to the fully equilibrated mixture of the three species (**i**). In view of the very complex trade-offs between the mechanisms and constraints, the following narrative should be regarded as a plausible interpretation of the qualitative features of this sequence rather than a rigorous account. Hord and Pucc start to invade in areas of highest resource densities (**b**). The competitively superior Hord thereby pushes Pucc into less habitable regions within the high-resource area they both occupy. This effect becomes more pronounced and also impacts Poa visibly as *N*_Hord_ and *N*_Pucc_ increase (**c**). Sharp vegetation zone boundaries start to appear and intensify for all species (**d** and **e**). The narrow high-density niches where Poa aggregates in **e** are almost vanquished in **f**, with Hord and Pucc filling the gap. Somewhat counterintuitively, two enclaves in the bottom right quadrant permit high density of Poa in **g**, despite the low resource concentration. Evidently, the minimisation of the total energy favours letting a part of Poa’s fixed (excessive) abundance 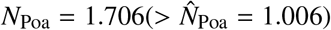 aggregate in a less preferable habitat as opposed to Poa bearing the cost of competition with Hord and Pucc in the high-resource areas. These two refuges disappear in **h** due to (i) increased competition with Hord and Pucc, and (ii) smaller *N*_Poa_. Overall, however, **g** already exhibits the main features of the final (smooth) equilibrium in **i**, with **h** depicting an intermediate stage.

**Extended Data Fig. 3.**
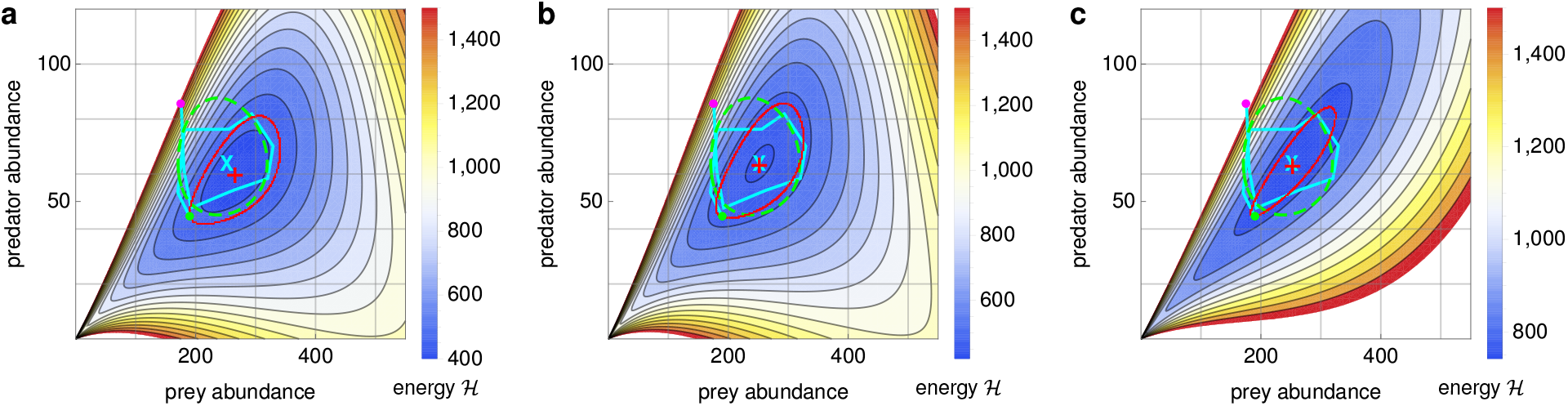
A suitable fitting procedure for Veilleux’s most stable predator–prey cycle. We illustrate the DFTe hypersurface 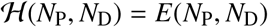 for equation (30) applied to the experimental setup of Supplementary Fig. 5a (the last fully visible cycle of the most stable predator–prey time series reported in Refs. [38, 39]), with parameters extracted from [39], see Supplementary Tab. 1. For a given DFTe equilibrium 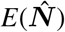 (red cross) we draw an equipotential line 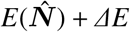 (red line) on 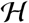, with *ΔE* such that the amplitudes of the actual data cycle (cyan line) around the average abundances ***N***^∗^ (cyan cross) roughly match. The equipotential lines drawn here and in Supplementary Fig. 6 are meant to test the predictive power of the DFTe hypersurface away from its minimum. The magenta dot marks the first minimum of P’s measured cycle, and the green dot marks the second minimum of D’s cycle. **a**, 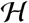 provides a reasonably accurate platform for the dynamics, with *γ* as the sole parameter for fitting 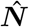 to ***N***^∗^. **b**, With two fit parameters (*γ*,*ν*_PD_), we obtain an even more satisfactory match between model and experiment, cf. Fig. 5. **c**, As one consistency check, we replace the common-sense choice of parasitism with amensalism and obtain a qualitatively different hypersurface: Comparing with **b**, we also find the equilibrium close to ***N***^∗^, but the DFTe trajectory is squeezed to such an extent that parasitism has to be regarded superior for modelling the predator–prey interaction mechanism. Moreover, only the hypersurface for parasitism permits noncyclic trajectories, which terminate as *N_D_* vanishes, for strong enough ‘excitations’ *ΔE*, that is, the predator population can collapse at high enough Cerophyl concentration while the prey survives, which is in line with experimental observations [39].

**Extended Data Tab. 3.** Synthetic community: Interactions, resource consumption, and dispersal. Each of the inter-species bipartite interactions, indicated in Fig. 6 with arrow heads that identify the primary direction of influence, is introduced with contact-type interaction kernel 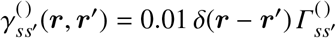. The matrices 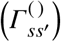 encode commensalism/amensalism (*a*), repulsion (*r*), mutualism (*m*), asymmetric competition (*c*), and parasitic interaction (*p*) that accompany predator–prey relations, see Extended Data Tab. 1. For example, 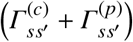 favours *s* = 4 over *s*′ = 1 and lets *s* = 3 benefit at the expense of its prey *s* = 1. To model a subset of species and resources, we simply extract the according submatrices of the matrices shown here. The first three rows of the resource requirements (*ν_ks_*) represent the prey species *s* = 1 and *s* = {2, 3}, consumed by the predators *s* = 3 and *s* = 7, respectively. In declaring the prey densities of Deer *ρ*_2_ =]*n*_2_[and Pig *ρ*_3_ =]*n*_3_[as separate resources for the Cat, we stipulate its reliance on each, cf. equation (7). If either prey could maintain the Cat, the combined resource density *n*_2_ + *ω n*_3_ were to replace *ρ*_2_ = *n*_2_ and *ρ*_3_ = *n*_3_, with a ratio *ω* to be specified. The rows 4–6 of (*ν_ks_*) identify the species’ requirements of the fixed equilibrium resource distributions *ρ*_4–6_ depicted in Extended Data Fig. 4. We ensure that resources can be exploited to a significant degree by choosing moderate with respect to the energy scale set by *ζ* = 1 in equation (7) dispersal pressures ***τ*** = 0.01 × (1, 2, 2, 0.3, 1, 1, 0.1) that penalise large local population densities only slightly.

| $(\Gamma_{ss'}^{(a)})$ | $(\Gamma_{ss'}^{(r)} + \Gamma_{ss'}^{(m)})$ | $(\Gamma_{ss'}^{(c)} + \Gamma_{ss'}^{(p)})$ | $(\nu_{ks})$ |
| --- | --- | --- | --- |
| $\begin{pmatrix} 0 & 0 & 0 & 0 & 0 & 0 & 0 \\ 0 & 0 & 0 & 0 & 0 & 0 & 0 \\ 0 & 0 & 0 & 0 & 0 & 0 & 0 \\ 0 & 0 & 0 & 0 & 0 & 0 & 0 \\ 0 & 0 & -1 & 0 & 0 & 1 & 0 \\ 0 & 0 & 0 & -1 & 0 & 0 & 0 \\ 0 & 0 & 0 & 0 & 0 & 0 & 0 \end{pmatrix}$ | $\begin{pmatrix} 0 & 0 & 0 & 0 & 0 & 1 & 0 \\ 0 & 0 & 0 & 0 & 0 & 0 & 0 \\ 0 & 0 & 0 & 0 & 0 & 0 & 0 \\ 0 & 0 & 0 & 0 & 0 & 0 & 0 \\ -1 & -1 & 0 & 0 & 0 & 0 & 0 \\ 0 & 0 & 0 & 0 & 0 & 0 & 0 \\ 0 & 0 & 0 & 0 & 0 & 0 & 0 \end{pmatrix}$ | $\begin{pmatrix} 0 & 0 & 0 & 0 & 0 & 0 & 0 \\ 0 & 0 & 0 & 0 & 0 & 0 & 0 \\ 1/2 & 0 & 0 & 0 & 0 & 0 & 0 \\ 1/2 & 0 & 0 & 0 & 0 & 0 & 0 \\ 0 & 0 & 0 & 0 & 0 & 0 & 0 \\ 0 & 0 & 0 & 0 & 0 & 0 & 0 \\ 0 & 1/2 & 1/2 & 0 & 0 & 0 & 0 \end{pmatrix}$ | $\begin{pmatrix} 0 & 0 & 1 & 0 & 0 & 0 & 0 \\ 0 & 0 & 0 & 0 & 0 & 0 & 5 \\ 0 & 0 & 0 & 0 & 0 & 0 & 1 \\ 1 & 1 & 1 & 1 & 1 & 1 & 0 \\ 0 & 0 & 0 & 0 & 1 & 1 & 0 \\ 1 & 1 & 1 & 1 & 1 & 1 & 0 \end{pmatrix}$ |

**Extended Data Fig. 4.**
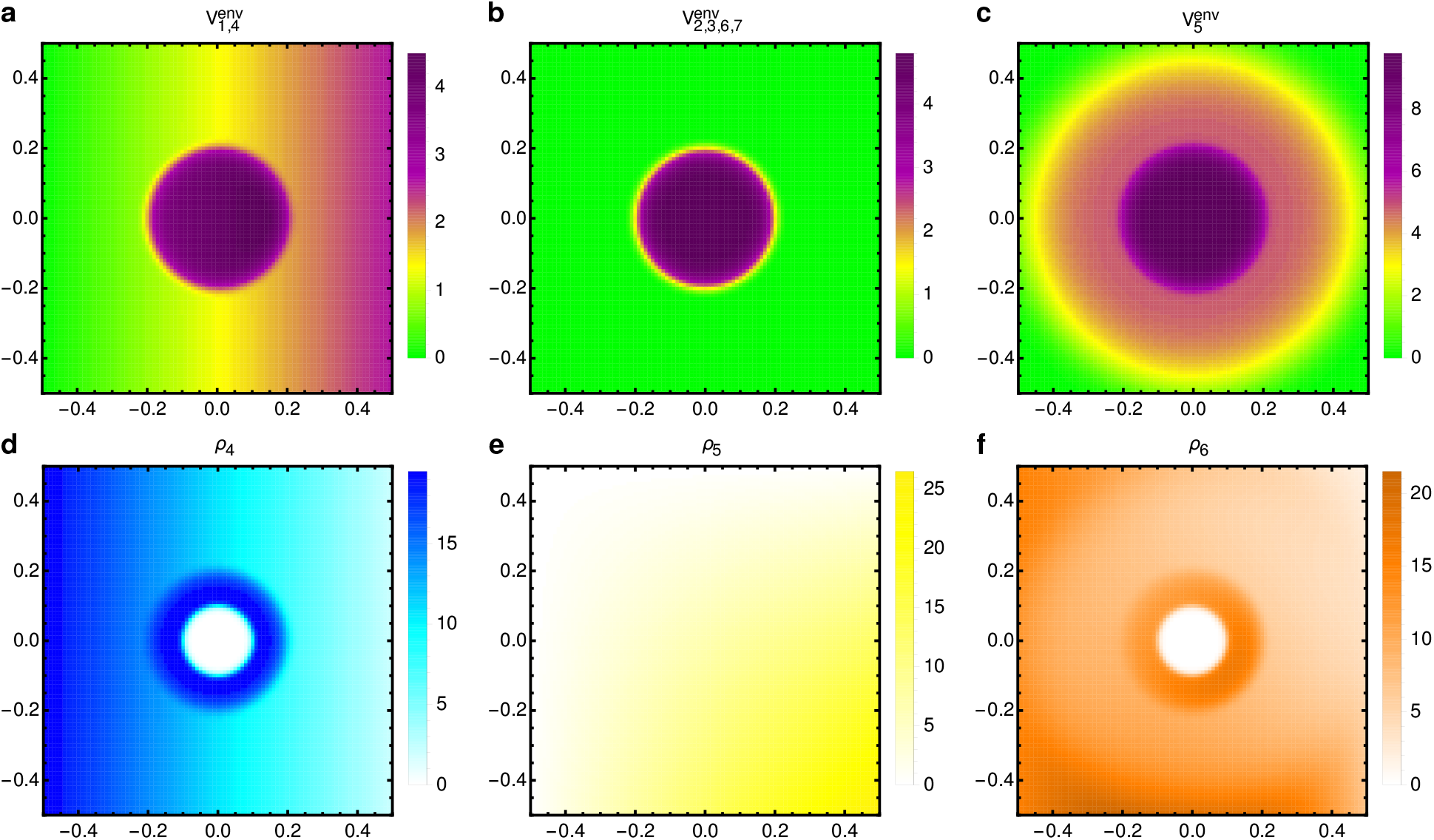
Synthetic community: Environments and resources. **a**–**c**, The species-specific environments 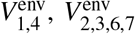, and 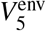 (all scaled with a factor of 100), from favourable (green) to hostile (black). **d**–**f**, The resource distributions *ρ*_4_ (‘water’) and *ρ*_5_ (‘sunlight’) are correlated with *ρ*_6_ (‘generic nutrient’). Choosing *σ* = −4 in equation (31), we assign the nonlimiting resources a minor but nonzero role relative to the limiting resources, see equation (8). We declare that the Trees’ environmental strain 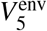 represents the deforestation stress, which is most detrimental in the central region. The central disk-shaped region is void of accessible resources *ρ*_4,6_, for example, due to human presence.

**Extended Data Fig. 5.**
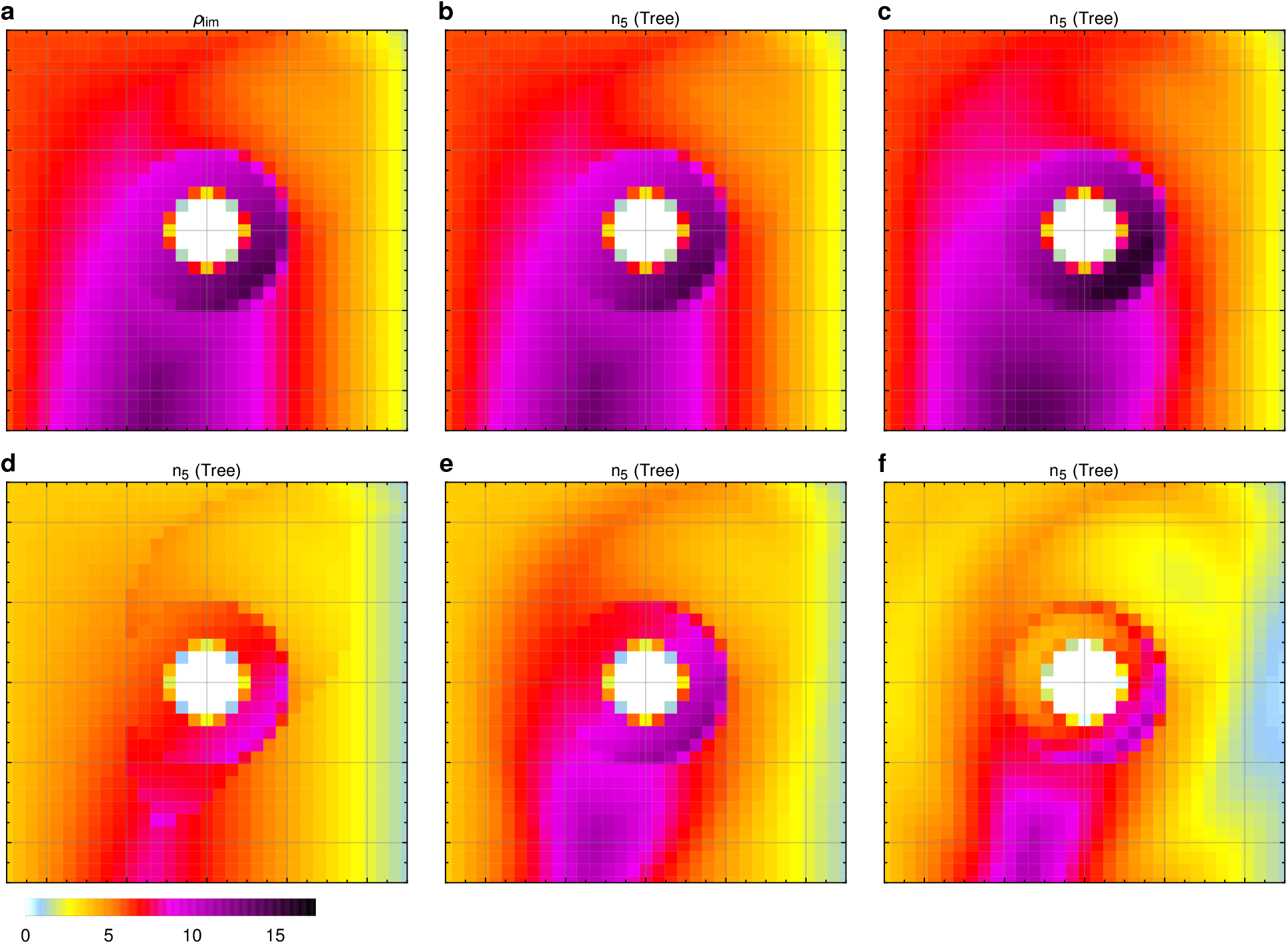
Synthetic community: Parameterisation of the Tree. We illuminate the interplay between dispersal, environment, and resources by modelling a single species (Tree, *s* = 5) with exclusive access to all resources. The color code applies to all panels. **a**, The locally limiting resource density 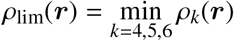 for the Tree, which requires equal amounts of *ρ*_4,5,6_. **b**, The equilibrium density distribution *n*_5_, with *σ* = −1000, *τ*_5_ = 10^−8^, and uniform environment (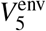 is replaced with zero), closely aligns with *ρ*_lim_ since (i) the dispersal pressure is essentially zero and (ii) the nonlimiting resources have negligible effect. **c**, The visible disparity to **b**is due to the smearing effect of the nonlimiting resources, triggered by setting *σ* = −4, which relaxes the constraints imposed by the limiting resources and permits higher overall consumption. **d**, Increasing *τ*_5_ to 0.01 while keeping *σ* = −1000, we reveal the adverse effect of the dispersal pressure from local conspecifics. **e**, The density for our generic choices *τ*_5_ = 0.01 and *σ* = −4, where the dispersal pressure is counteracted by the density-increasing effect of the nonlimiting resources. **f**, The introduced environment 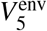 (see Extended Data Fig. 4) expels the Tree from hostile regions — to an extent that is determined by balancing (through the minimisation of the DFTe energy) the according environmental pressure against (i) the dispersal pressure and (ii) the conspecific resource competition.

**Extended Data Fig. 6.**
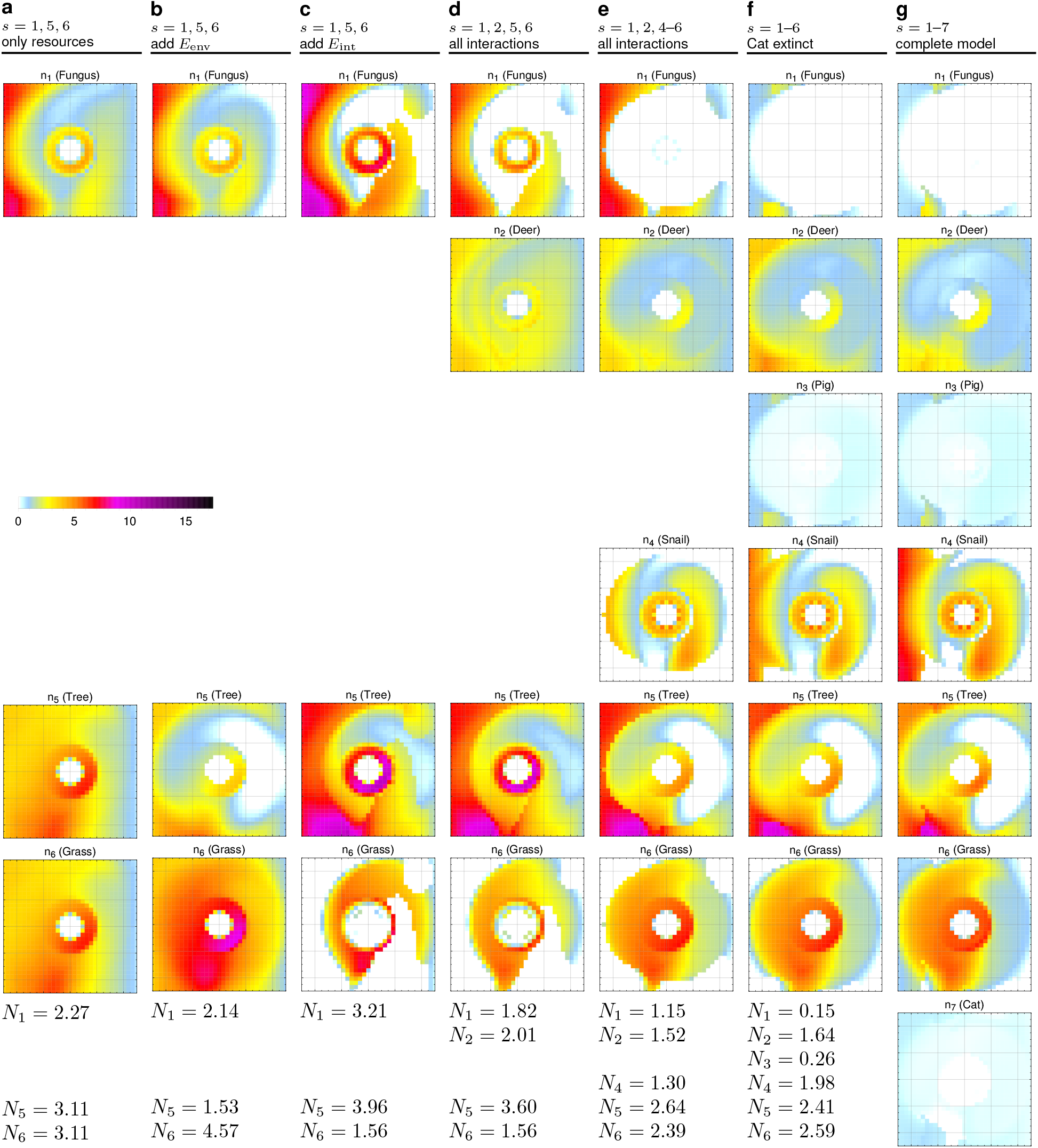
Building the synthetic community bottom-up. Each column (**a**–**g**; shared colour code) specifies the involved species and the energy components that are successively added to *E*_dis_; see Supplementary Notes for a detailed interpretation of **a**–**g**. The transition from **g**, with abundances ***N*** = {0.13, 1.26, 0.29, 2.53, 2.04, 2.69, 0.19}, to **f** amounts to an artificial extermination of the apex predator (Cat), that is, the absence of the Cat in **f** is not the result of competitive exclusion but of external intervention. The according relative redistributions of *s* = 1–6 are spatially resolved in Fig. 6b. The main insights are (i) the decline of Grass and Snail populations, (ii) increased abundances of their heterospecifics through feedback loops, (iii) knowledge on how the inter-species mechanisms, coupled with the environmental constraints, lead to these results, and (iv) which specific actions to take for expanding or helping reintroduce the Cat population (see Extended Data Fig. 7).

**Extended Data Fig. 7.**
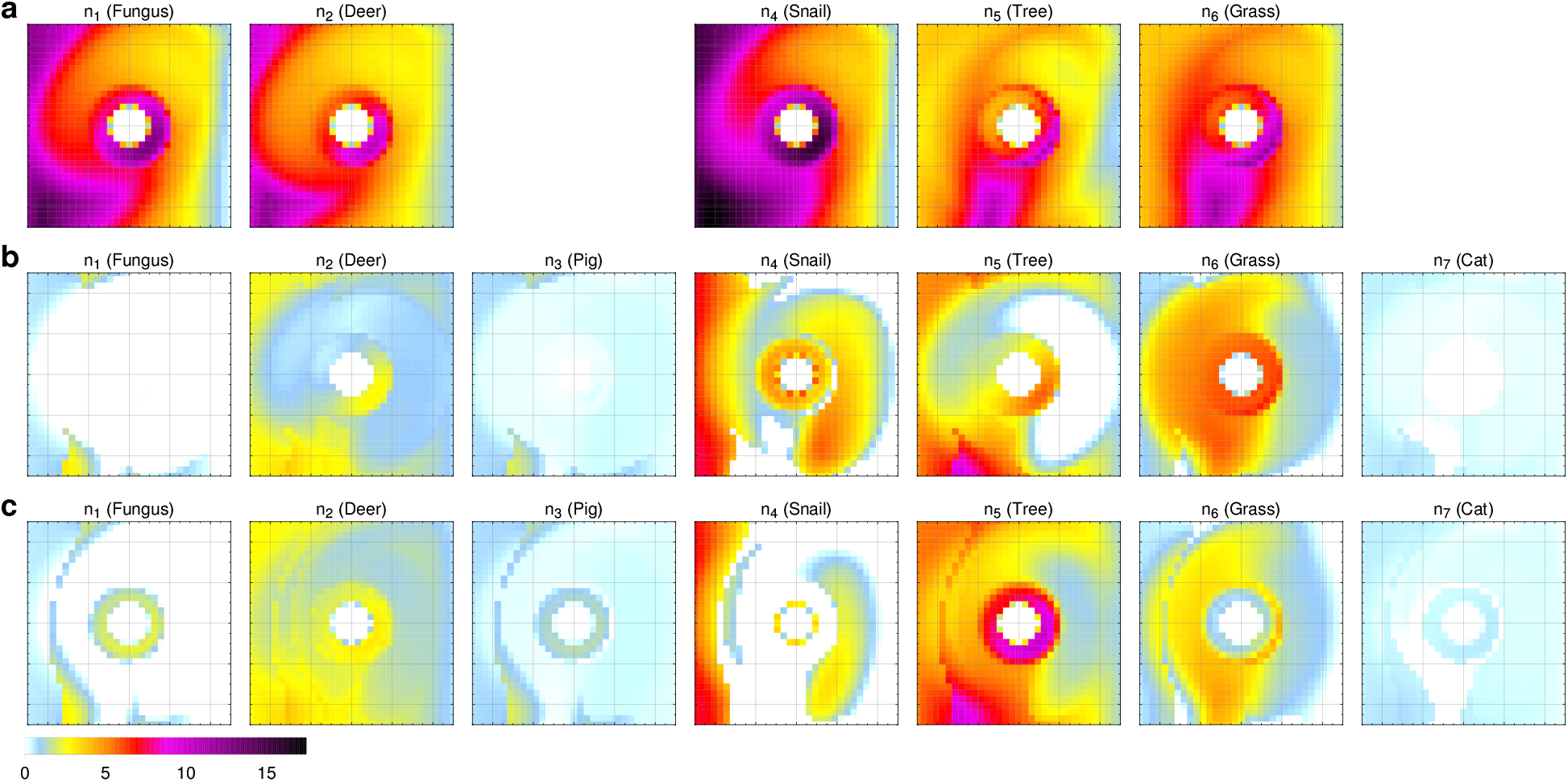
Synthetic community: Species in isolation, at equilibrium, and in modified environments. Interactions can considerably transform species distributions and decorrelate them from the resource distributions. The colour code applies to all panels. **a**, Distributions of the species *s* ∈ {1, 2, 4–6} in isolation, that is, in absence of all *s*′ ≠ *s* and solely constrained by the resource distributions and the *s*-specific environment. We disregard the predators Pig and Cat, which rely on prey resources and thus inherently require inter-species interactions. While Snail and Fungus perceive the same environment and require the same resources, the Snail is a more efficient consumer due to its lower dispersal pressure, which results in higher densities. The Deer experiences a favourable environment except in the central region, but its high dispersal pressure leads to lower densities overall. The Tree and Grass only differ in their environment, see Extended Data Tab. 3 and Extended Data Fig. 4. **b**, Distributions (identical to column **g** of Extended Data Fig. 6) for the complete interacting community, including predation. **c**, We expand the Cat population by cutting the deforestation stress for the Tree in half see Supplementary equation (S18). Evidently, this measure not only increases the Cat abundance from *N*_7_ = 0.19 to *N*_7_ = 0.26, but affects the entire community significantly. The relative difference between **b** and **c** is depicted in Fig. 6c.

## SUPPLEMENTARY INFORMATION

### Supplementary Notes

#### Notes on the DFTe energy functional

By minimising the energy *E* = *E*[***n***, ***μ***](***N***) in equation (9), we implicitly state that the ecological system prefers realisations of lower energy. At its core, the modular decomposition of *E* is one of notation and not based on fundamentally disparate entities. For example, the dispersal energy in equation (4) can be thought of as an internal interaction energy, and the environmental energy can be viewed as an interaction between the system constituents and the environment. In this language we would term *E* a pure interaction energy. Splitting *E* into the parts of equation (9) merely serves the purpose of introducing and building on intuitive concepts when we determine the functional form of *E*. Furthermore, only relative differences to a reference energy are of consequence. For example, we may shift and/or rescale *E* such that its minimum equals one. Since (i) energy can be rescaled and (ii) quantities of energy and density are the only dimensionful objects that enter *E*, the only relevant unit in DFTe is length. In principle, the energy in equation (9) could be expressed in terms of any energy unit (e.g., Joule), but this would be (i) a tedious and ambiguous enterprise in practice and (ii) of no relevance to our analysis of ecosystems with DFTe. The parameterised components of *E* are not fitted to energetic data. We rather fit the *minimisers* of *E* to our only objects of interest, namely density distributions and abundances.

A ubiquitous difficulty in describing many different species that interact within a complex environment is to find the proper trade-offs among given mechanisms and constraints. And this agenda gets more involved as the complexity increases. One strength of DFT-type approaches is that the complexity of the underlying functional *E* does not matter much as the trade-offs between the components are found automatically in the minimisation process, while the building blocks of *E* are parameterised independently from less complex subsystems. Although the numerical minimisation of *E*[***n***, ***μ***](***N***) for any given system can be technically involved, the greater challenge for DFT usually lies in determining the explicit forms of the required functionals. Here, we have identified an energy functional *E*[***n***, ***μ***](***N***) for ecology that lends itself to straightforward interpretations and quantitatively describes a wide range of ecological systems, despite its structural simplicity and with very few fitting parameters.

With a specific system at hand, we retrieve the appropriate parameter values in equation (9) from actual data. Once these values are identified and validated, we can predict the species densities in new situations. Most importantly, we can predict the behaviour of complex systems by minimising *E*[***n***, ***μ***](***N***) using the parameters obtained for less complex subsystems. We can also rule out interaction mechanisms that yield poor fits to data. That said, equation (9) is likely not the last word on predicting species distributions in ecology. But we can and do show consistency of equation (9) with experimental and field data across a variety of taxa and situations. In the course of studying specific experimental systems, we have justified all components of equation (9). While DFTe predictions based on less controlled field data have to be the final criterion for judging the universal applicability of equation (9) to ecology, here we focus on establishing the structure of DFTe from relatively unambiguous laboratory data and established synthetic data.

#### Notes on the resource energy

The DFTe energy in equation (9) balances the cost due to dispersal, environment, and interactions with the cost *E*_Res_ of foregone resources. The latter are replenished in equilibrium and are usually consumed only in part due to the multiple constraints in a complex ecosystem. The rate of consumption is irrelevant since *ρ_k_*(***r***) is replenished at the same rate in equilibrium. For example, a 50-hectare pasture that grows biomass at 20 kg/(ha day) sustains an equilibrium abundance of 50 cows (each requiring 20 kg of grass daily), such that we may choose *ν* = 20 and *ρ* = 1000 in equation (7). Expressing *E*_Res_ as the sum of

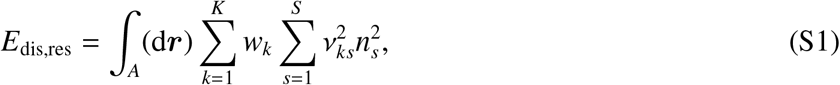

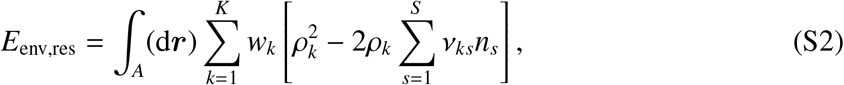

and

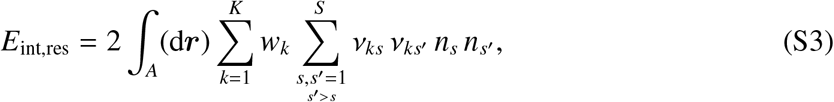

we reveal the rich interpretations of the simple expression in equation (7). *E*_dis,res_ encodes the intraspecies pressure in consuming resources, akin to the dispersal energy in equation (4). The negative term 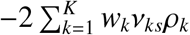 in *E*_env,res_ modifies the environment 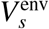 favourably and balances against the positive *E*_dis,res_ and *E*_int,res_. The constant offset produced by 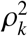 is irrelevant to the minimiser of *E* (prior to the addition of other energy components of equation (9)). *E*_int,res_ presents inter-species pressure mediated by resources (just like *E*_dis,res_ does for conspecifics) and comes in the form of a repulsion symmetric in *s* and *s*′, see Extended Data Tab. 1. It is a special case of the more general interaction energy

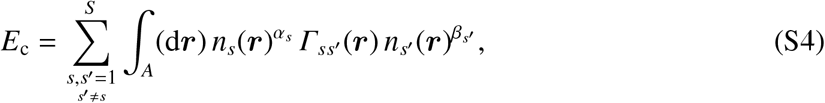

which is itself a special case of equation (6) with *γ_ss′_* (***r***, ***r***′) = *δ*(***r*** − ***r′***) *Γ_ss′_* (***r***). In addition to the resource-based pressure, internal pressure may come from other intra-species mechanisms, such as competition for mates or territory, and can be encoded in *τ_s_*, which lets the equilibrium value of 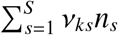 in equation (7) deviate from its a priori equilibrium value *ρ_k_*.

#### An analytically solvable minimal example with resource competition

In the following, we illustrate the interplay of resources and interactions with the help of two minimal, synthetic examples. We consider uniform systems of two species that tap into resources *R_k_* = *Aρ_k_* = 3 in an area *A* = 1 and exhibit amensalism of interaction strength *Γ* = *Γ*_12_, which puts species 2 (‘s2’) at a disadvantage relative to species 1 (‘s1’), see Extended Data Tab. 1.

First, employing a single resource and specifying *ν*_11_ = 2 while keeping *ν*_12_ > 0 variable, that is, 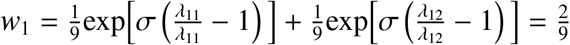, we minimise the total energy

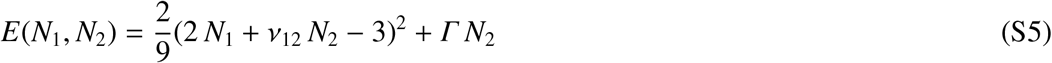

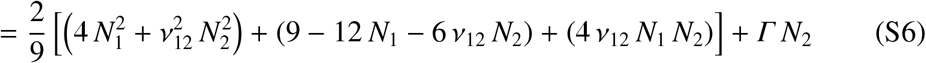

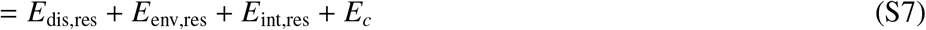

for the admissible *N_s_* ≥ 0. For *Γ* = 0, one might expect that an increase of *ν*_12_, which increases the internal pressure for s2, results in an expansion of s1. However, the dispersal energies are then exactly balanced with the increased ‘resource interaction’ *E*_int,res_ between both species and the more favourable ‘resource environment’ 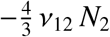 for s2. Indeed, the global minima of equation (S5) are 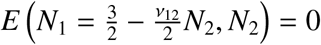 for *Γ* = 0, with *N*_2_ restricted by 0 < *N*_2_ < 3*/ν*_12_. Coexistence of two species sustained by one resource is therefore possible for a continuum of abundances. However, for any *Γ* > 0 the global minimum 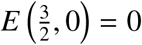 is unique, which means that even an infinitesimal competitive interaction settles the ambiguity in favour of the competitively superior s1.

The two species of our second toy system require two resources (*ν*_11_ = 2, *ν*_12_ = *ν*_21_ = 1, variable *ν*_22_ > 0). Apart from the numerical values of the parameters, the according energy functional is the one used to describe Tilman’s two-species competition experiments that led us to Fig. 3. With the carrying capacities

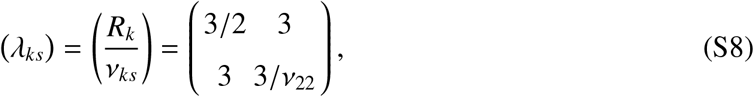

the limiting resource for s1 is always *R*_1_, while s2 is limited by *R*_2_ (*R*_1_) for *ν*_22_ > 1 (*ν*_22_ < 1). We fix the weights *w_k_* of equation (8) by *σ* = −4 and then have to minimise

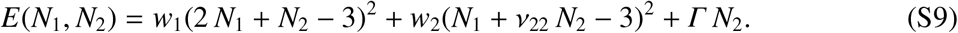

**Supplementary Fig. 1.**
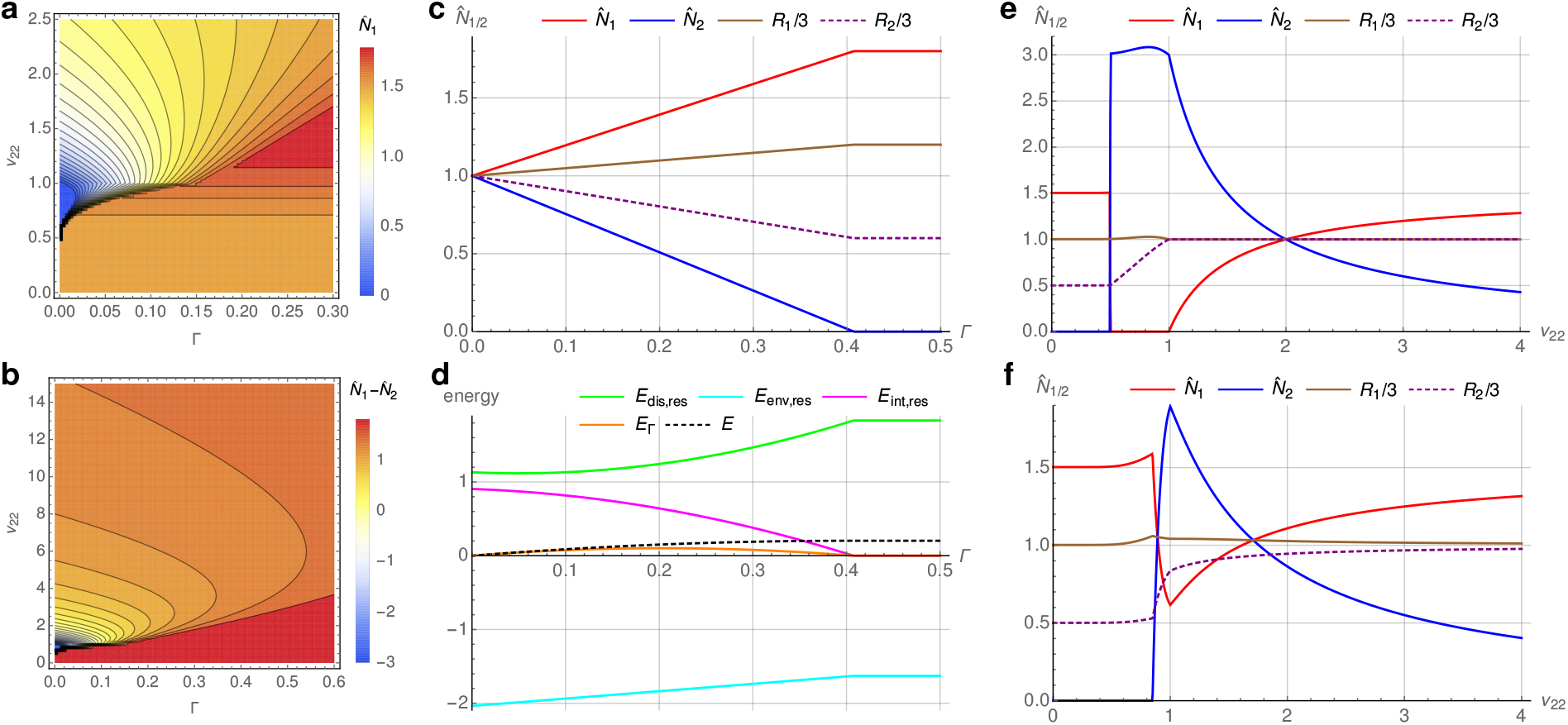
DFTe predictions for a synthetic minimal community. Tuning interaction strength *Γ* and resource requirement *ν*_22_ of the two-species s1ystem defined through equation (S9), we explore the trade-off between resource- and interaction energy, which leads to a rich variety of equilibrium states. Panel **a**shows the abundance 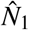 of s1, which is (generically) superior to s2, except in a small region of the parameter space where the interaction strength *Γ* is small enough and where the resource requirement *ν*_22_ of s2 falls into a suitable window. The difference 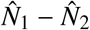 in panel **b**hints at the asymptotic behaviour (i.e., competitive exclusion of s2) for large *Γ* and *ν*_22_. Panels **c** and **d** show how the departure from the symmetric situation (*Γ* = 0 and *ν*_22_ = 2, resulting in 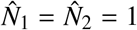) is accompanied by automatically determined trade-offs between the interaction energy proper *E_Γ_* and the resource-based energy components *E*_dis,res_, *E*_env,res_, and *E*_int,res_. We show the phases and phase transitions that abundances and resources undergo as *ν*_22_ varies for constant *Γ* = 0 (**e**) and *Γ* = 1/18 (**f**).

The global minimisers 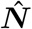 of equation (S9) are shown in Supplementary Fig. 1a/b, which demonstrates that already such a simple system of two species exhibits gradual suppression of either species and competitive exclusion of one species sharply turning into competitive exclusion of the other species. Supplementary Fig. 1c depicts the gradual suppression of s2 by s1 as *Γ* increases at fixed *ν*_22_ = 2, until s2 is competitively excluded beyond the critical value 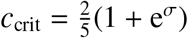. As *E*_Res_ has to be balanced against *E_Γ_* for *Γ* > 0 by permitting an energy cost that comes with foregone (*R*_2_) and overconsumed (*R*_1_) resources, the precise consumption of all available resources cannot be maintained. Supplementary Fig. 1d shows the energy components for the situation of Supplementary Fig. 1c. *E*_int,res_ is largest for *Γ* = 0, where *N*_1_ = *N*_2_, and shrinks as the abundances increasingly deviate from the symmetric situation, until s2 is excluded. These drastic changes are driven by an amensalism energy *E_c_* that is small throughout in comparison to the other energy components. Although *N*_2_ vanishes beyond *c*_crit_, where *E_c_* = 0, amensalism is still a property of the system: s2 cannot invade if *Γ* > *Γ*_crit_ since its introduction would cost too much interaction energy *E_c_*. This energetic condition, which emerges in the presence of amensalism during the energy minimisation, then shows up in the changed *E*_dis,res_ and *E*_env,res_ (compared with their values at *Γ* = 0), which add up to a finite positive value of the total energy. Supplementary Fig. 1e for the noninteracting case (*Γ* = 0) exhibits the basic phases of the system. Both species are on equal footing in the symmetric situation for *ν*_22_ = 2, while *ν*_22_ > 2 (*ν*_22_ < 2) increases (decreases) the intra-species pressure for s2 compared to s1. Increasing *ν*_22_ beyond *ν*_22_ = 2 therefore excludes s2 gradually. In a nonuniform environment an increased intra-species pressure for one species would push its density distribution into previously (too hostile) unoccupied territory, see also Extended Data Fig. 1. In a uniform environment its only effect is an increased cost due to increased intra-specific repulsion, such that the species as a whole cannot consume as many resources. The so released resources become available to other species. Conversely, a decreasing *ν*_22_ gradually excludes s1, until it vanishes for *ν*_22_ = 1. For all *ν*_22_ > 1 the two species are limited by different resources. As *ν*_22_ decreases below one, *R*_1_ becomes the limiting resource also for s2 (such that *N*_2_ = 3), with an increasing amount of unutilised *R*_2_ as *ν*_22_ approaches 1/2 (note that not the resource energy has to be minimised, but the total energy, such that lowering the total energy through overconsumption is permitted, which is why s2 exceeds *N*_2_ = 3 for 1/2 < *ν*_22_ < 1). The cost of foregone *R*_2_ is then so high that s1 enters the scene again — and in doing so, expels s2 completely. Already a simple system of two species (that only interact via resource consumption) thus exhibits gradual exclusion of either species (1 < *ν*_22_ < 2 and *ν*_22_ > 2), and competitive exclusion of s1 (for 1/2 < *ν*_22_ < 1) sharply turning into competitive exclusion of s2 (for 0 < *ν*_22_ < 1/2). Supplementary Fig. 1f for finite amensalistic interaction strength *Γ* = 1/18 shows the same qualitative picture as Supplementary Fig. 1e, but s2 dominates only for 0.9 ≾ *ν*_22_ ≾ 1.7 instead of 1/2 < *ν*_22_ < 2, and s1 is never fully excluded. Finally, we illuminate the role of *σ* in the weights of equation (8) by observing that *σ* → −∞ yields (*w*_1_*, w*_2_) = (1/9, 1/9) for *ν*_22_ > 1. Both resources are equally important, and the two species coexist with unique abundances for each value of *ν*_22_. For *ν*_22_ < 1, both species are limited by *R*_1_, such that (*w*_1_*, w*_2_) = (2/9, 0) reduces the system to our first example above (with *ν*_12_ = 1). In contrast, a finite *σ* implies finite weights for both resources and all *ν*_22_, resulting in the unique abundances shown in Supplementary Fig. 1e/f. Our conclusion for this minimal example of two species therefore is that (i) the advantage of s1 over s2 due to amensalism is an advantage for all resource requirements *ν*_22_ — relative to the noninteracting case — and (ii) either species can suppress the other depending on *ν*_22_ (or, alternatively, the provided resource combination).

#### Notes on Fig. 2

In Ref. [25] (see also [51]), Méndez-Valderrama *et al.* set up a ‘density-functional fluctuation theory’ (DFFT) and obtain remarkably accurate density distributions of a single species (*Drosophila melanogaster*) in a heterogeneous environment (holding chambers with one side exposed to a heat source). They employ the grand-canonical ensemble of statistical physics with an energy functional whose system-specific parameters are extracted directly from the data: While they assume an environmental energy functional of the form in equation (5) to capture the preferential positions of fruit flies subjected to a temperature gradient in the chambers, the intra-specific interaction functional remains unspecified. This approach is particularly useful if nothing is known about those interactions or if no functional form should be assumed. Its applicability to more complex situations remains to be studied, though. Most importantly, resources are not considered since a fixed number of fruit flies is constrained to the chambers for a limited time. This situation represents the fundamental building block of our DFTe framework: Calculating the density distributions for fixed abundances, whatever the energy functional. In contrast to DFFT, we have to specify an interaction functional and explicitly parameterise an environment. This can be seen as a disadvantage since we impose mechanisms and structure on the ecosystem model. But it also means that we are able to discriminate between mechanisms, can dismiss those that are inconsistent with the data, and thereby gain insight that is directly interpretable.

We extract a smooth and monotonic reference density 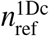 from the experimental data points from the quasi-1D chamber (with area *A* = 10 × 0.8 cm^2^ and 24 × 2 bins) of Ref. [25] by averaging pairs of the 48 bins in descending order. The cubically interpolated result is shown as the red dotted line in Extended Data Fig. 1 and resembles an inverted parabola, such that the density expression in equation (2) yields a first approximation of 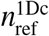 for *V*(***r***) = *V*^env^(***r***) ∝ (*x* + *x*_0_)^2^ and *x*_0_ = 5 cm.

We assess the quality of predictions vs data with the least-squares correlation measure

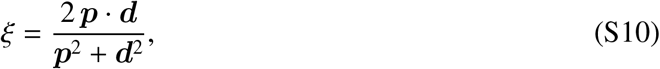

illustrated in the table below. We also note that uniformly drawn unit vectors ***p*** with positive entries in three dimensions yield the expectation value 〈*ξ*〉 = 1/2 for any given unit vector ***d***.

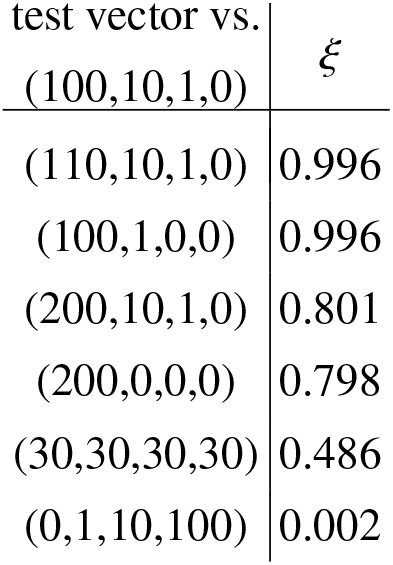

#### Notes on Fig. 3

We find a valuable test system for DFTe in Tilman’s seminal study of resource competition (*R*^∗^-theory) among four diatoms of Lake Michigan [1]. While the experimental data of abundances in Ref. [1] are limited to two species competing over two resources (SiO_2_,PO_4_) for 30 days (validating the *R*^∗^-predictions reasonably well), *R*^∗^-theory is applicable to any number of species and resources over any period of time. Although both data and predictions in Fig. 6 of Ref. [1] seemingly approach equilibrium abundances of coexisting F and A, the actual *R*^∗^- equilibrium from Tilman’s fitted model (approached as the observation time tends to infinity) amounts to competitive exclusion of A and F, respectively, for two of the three employed resource combinations, see Extended Data Tab. 2. In fact, besides cases with involvement of the inferior T, for which both data and *R*^∗^-theory suggest exclusion of T under all circumstances, five out of the nine remaining cases are not even remotely (*R*^∗^-)equilibrated after 30 days. Since we want to test the equilibrium predictions of DFTe, we thus refrain from benchmarking against the actual (transient) data and rather aim at reproducing the equilibrium abundances that follow from *R*^∗^- theory. Strictly speaking, we are not dealing with a two-dimensional system here, but one may simply think of the third direction integrated out. Without loss of generality we fix the unit of length via *A* = 1, which amounts to 1 mL suspension by our definition. The unit of energy is then implicitly fixed via *ζ* = 1 in equation (7).

The energy function *E*(***N***) in equation (28) requires input from monoculture data only. The *R*^∗^-values

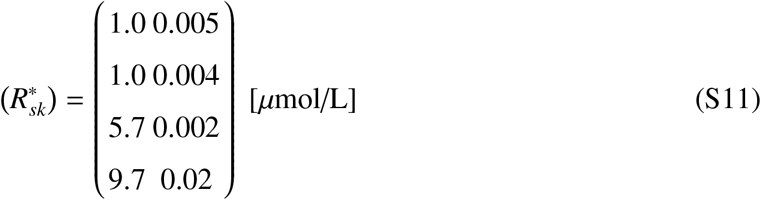

and nutrient requirements

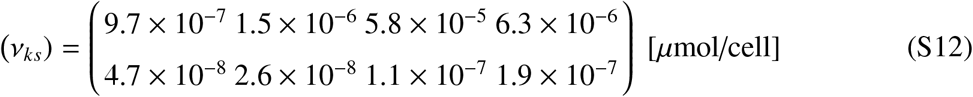

that enter equation (28) are taken from Tab. 1 of Ref. [1]. The two free parameters of *E*(***N***) (*γ* and *κ* in the interaction term) are fitted exclusively to all two-species subsets of (F|A|S|T) subjected to the reference resource combinations 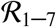, with an average of *ξ* ≈ 0.992 for these 42 cases. The best fit (*γ* = 8 × 10^−8^ and *κ* = 8) maximises *ξ* among all those parameters that yield the correct survivors (according to *R*^∗^-theory). Supplementary Fig. 2 shows a histogram for 500 randomly drawn resource combinations that yield correlations of *ξ* ≈ 1 between DFTe and *R*^∗^-theory, as discussed in Fig. 3. Complete correlation (*ξ* = 1) is not expected since the generic resource energy of DFTe is void of species-dependent threshold resource densities *R*^∗^, which are corner stones of Tilman’s resource competition theory, and thus permits complete consumption of all provided resources. Accordingly, small values of *ξ* (like for (F|A|S) subjected to 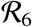, see Extended Data Tab. 2) are encountered for resource combinations close to the *R*^∗^-values. We emphasise, however, that the quantitative differences between DFTe and *R*^∗^-theory in such cases is largely accounted for by DFTe’s (over-)consumption of magnitude *R*^∗^. If we artificially reduce the *R*^∗^-values by a factor of 1000, *R*^∗^-theory indeed predicts (F|A|S) = (0|72|170) for 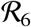. A corrective energy term that captures each species survival thresholds of resources densities 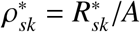 can be introduced, for example, through a species-specific environmental potential

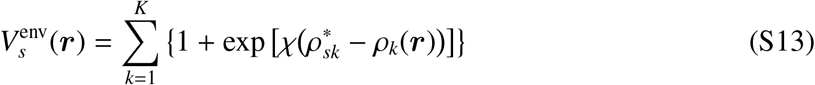

which exponentially penalises species *s* at ***r*** if any *ρ_k_*(***r***) approaches or falls below 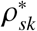. A priori, *χ* is an additional (likely large positive) fit parameter.

With *κ* = 8, the fitness proxies *g_s_* based on resource requirements per cell are virtually irrelevant compared to the fitness proxies *f_s_* that feature the *R*^∗^-values. They are not completely superfluous, though, since F and A both exhibit *R*^∗^ = 1 under SiO_2_ limitation, which would render both species noninteracting [in the sense of equation (25)] if it were not for their differing resource requirements. Of course, this is a fine-tuning artefact of numerically identical *R*^∗^-values — that is, equation (25) is likely independent of *ν* in realistic communities. Furthermore, it would be surprising if equation (25) were the only possible choice for the interaction kernel, and it is quite likely that many different energy functionals that encode different mechanisms and input parameters are equally suited for reproducing the data targeted in this example. Only future DFTe studies that take into account additional data can reduce the space of possible functionals. Taking the cue from physics, we may hope that a universal functional will prevail and thereby provide deeper insights into the fundamental mechanisms and relations underlying ecological systems in general and microbial communities in particular.

**Supplementary Fig. 2.**
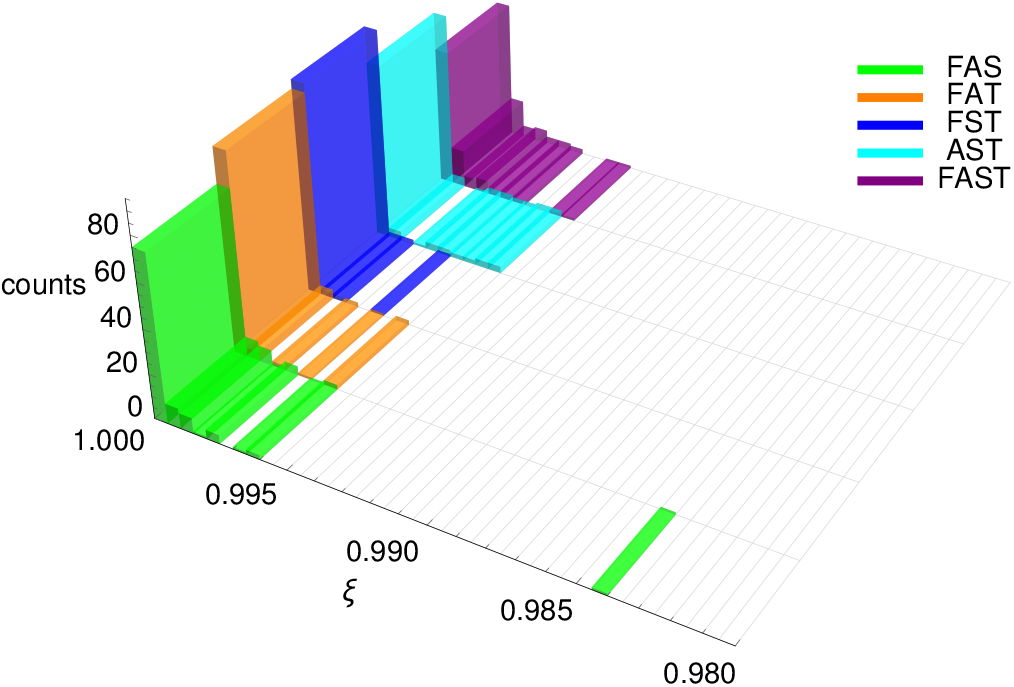
Summarising histogram of the results shown in Fig. 3. Comparing the DFTe predictions with the *R*^∗^-predictions for randomly chosen resource combinations [100 cases each for (F|A|S), (F|A|T), (F|S|T), (A|S|T), (F|A|S|T)] in the ranges [9.991 … 500] for SiO_2_ and [0.0206 … 20] for PO_4_, we find a near-perfect match in all cases. The lower limits of the resource ranges are 3% above the maximal *R*^∗^-values (9.7 SiO_2_ and 0.02 PO_4_ for T). The smallest overlap (*ξ* ≈ 0.984) between DFTe and *R*^∗^-theory among all our resource cases is (incidentally) encountered for one resource case of (F|A|S), see Extended Data Tab. 2.

#### Notes on Fig. 4

The purpose of this example is two-fold. Our first goal is to predict densities of species in a nonuniform environment using data of experiments carried out in a uniform setting. Second, we apply DFTe to data that come with very limited information on mechanisms and species properties — a situation that requires us to postulate a functional form of *E* (informed by the available data) and to fit many parameters of *E*.

Kenkel *et al.* in Ref. [36] report the response of three grasses [*Poa pratensis* (Poa), *Hordeum jubatum* (Hord), and *Puccinellia nuttalliana* (Pucc)] to eight salinity levels. All three plants perform best at lowest salinity when grown in monoculture, but competitive interactions change this pattern when grown in mixture, see Supplementary Fig. 3a, where we replicate the data of Ref. [36] for the above-ground biomass in grams, collected after 94 days of growth from pots of 10 cm diameter (corresponding to *A* = 1 in our units) that were seeded with 150 plants per species. All experiments were performed in triplets, with averages presented in Supplementary Fig. 3a for each of the eight NaCl concentrations from 0 to 14 g/L and for each species in both monoculture and mixture. The salinity level is constant in each experimental pot, such that the data points in Supplementary Fig. 3a are to be viewed as isolated. We assume that the data represent equilibria. The final abundances are bounded from above by the constant concentration of the nutrient medium (which includes the salt, but remains unspecified otherwise) and accessible pot area. Without loss of generality, resource *R_s_* limits species *s* with *ν_ss_* = 1, such that the monoculture abundance *N_s_* in Supplementary Fig. 3a equals the provided amount of resource *R_s_* (for *τ_s_* = 0). These limiting resources also put upper limits on the six off-diagonal resource requirements *ν_ks_* with *k* ≠ *s*. We use an interaction strength parameter *γ* factored into *γ_ss′_* = *γ* [*f_s_/ f_s′_* − 1]_+_ and the same fitness proxies *f_s_* for all three types of competitive bipartite interactions (amensalism, repulsion, and asymmetric interactions, see Extended Data Tab. 1). As Fig. 4 of Ref. [36] suggests, we fix the salinity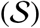-dependent fitness proxies *f_s_* as the fraction of above-ground biomass in mixture, approximated by the shifted Gaussians

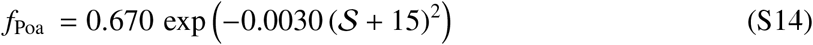

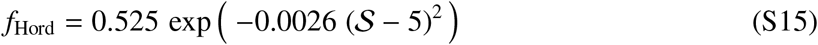

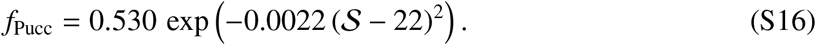

The dispersal encodes an intra-species repulsive interaction, which supplements the inter-species interactions. Finally, the free parameter *σ* in equation (8) allows us to fit the weights *w_k_* of the three resources. We employ the units of length and energy, implicitly defined by *A* = 1 in equation (29) and *ζ* = 1 for the resource energy in equation (7), respectively.

We now ask what happens if we connect the pots, such that the grasses are allowed to aggregate in their preferred habitat of low salinity at the expense of stronger competition. This question obviously emerges as we shift focus from the laboratory to realistic environments. We pursue an answer by first fitting the nine parameters of the DFTe energy functional in equation (29) to the constant-salinity data and then predict the density distributions of Poa, Hord, and Pucc in heterogeneous saline environments. We never find zonation at equilibrium, see Supplementary Fig. 3b–d and Supplementary Fig. 4, where the equilibrium density profiles turn out to be void of any discontinuities. This observation holds across the three basic types of competitive interactions and for all salinity landscapes considered, be it a linear (lin) salinity distribution or randomly (ran) chosen ones. However, a rich zoo of phases emerges as equilibrium is *approached*: Asymmetric interactions (in the general sense of one species benefiting at the expense of another, see Extended Data Tab. 1) yield zonation, competitive exclusion, and other discontinuities in out-of-equilibrium situations, see Extended Data Fig. 2 and Supplementary Fig. 3e.

**Supplementary Fig. 3.**
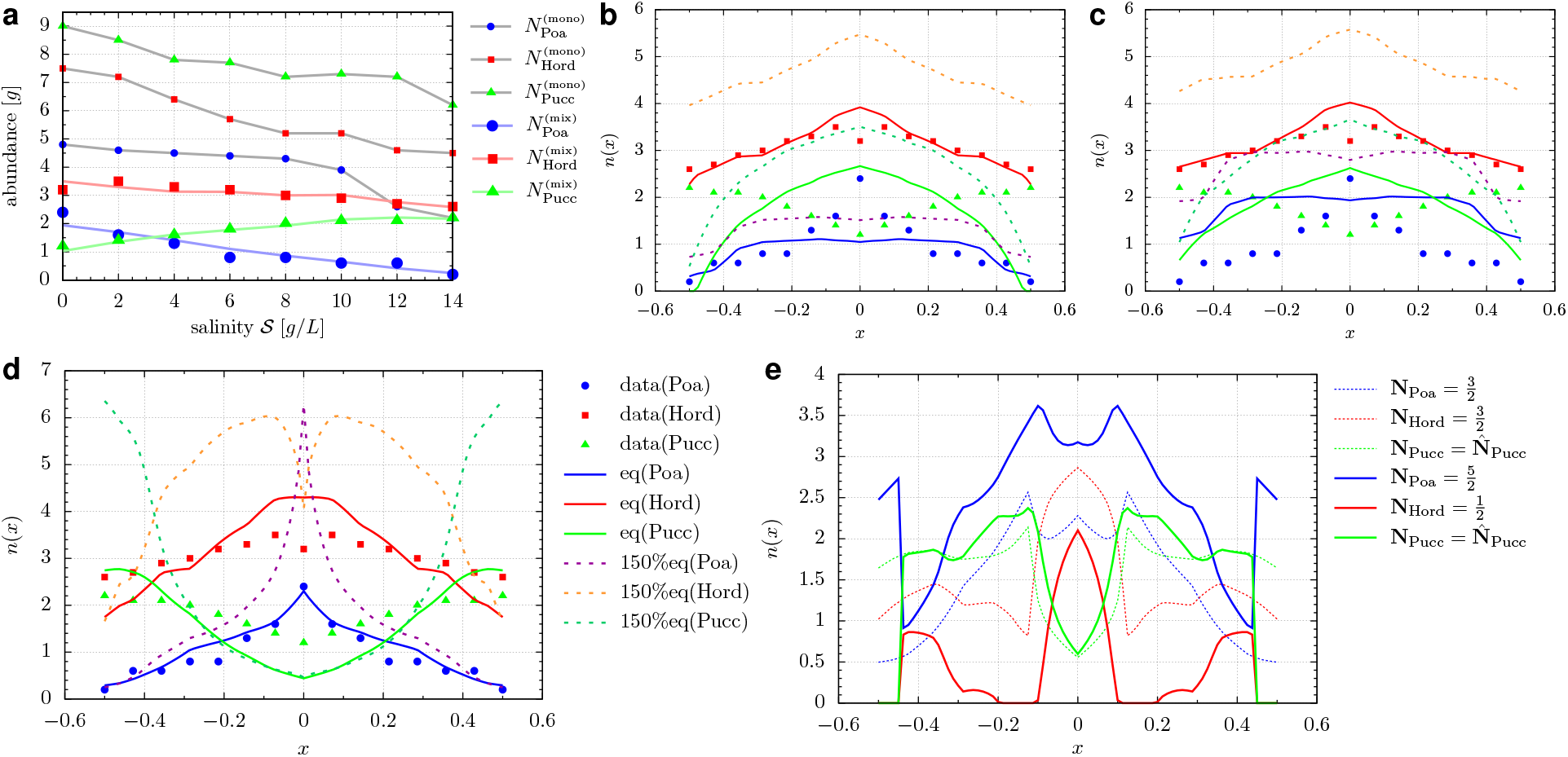
Fits to uniform salinity inform DFTe predictions for a salinity gradient. **a**, The experimentally observed equilibrium abundances (in grams) of Poa, Hord, and Pucc as a function of (constant) NaCl concentrations (in grams per litre). Salinity-dependent competition of some kind is at work since the mixture abundances *N*^(mix)^ are not simply rescaled monoculture abundances *N*^(mono)^. The grey lines through the monoculture data guide the eye. The colour lines follow from the best fits of the DFTe function in equation (29) with asymmetric interaction to the mixture abundances, see Supplementary Fig. 4. The fits for amensalism and repulsion are of similar quality. We use the resulting fitted DFTe parameters to predict the equilibrium density profiles from minimising the nonuniform version of equation (29) for a ‘V’-shaped linear salinity distribution 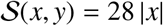 (constant in *y*-direction). The salinity 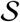 is mapped to the values of the resources *R*_1_, *R*_2_, and *R*_3_ by means of the functional relationship between 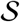 and the monoculture abundances of panel **a**. Obviously, the three resources are highly correlated. Panels **b** and **c** for amensalism and repulsion, respectively, show similar trends when compared with the uniform abundances: Hord is largely unaffected, Poa’s distribution flattened, and Pucc inverted in response to the salinity gradient. In particular the latter outcome contrasts with field observations, where Pucc is competitively displaced to areas of high salinity levels [52]. We recover the trends of these field observations when employing asymmetric interactions in panel **d**. Forcing the system into a nonequilibrium high-density state [‘150%eq(·)’] by fixing the abundances at 50% above the equilibrium 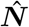 [‘eq(·)’], we reveal more clearly the habitat niches that each species prefers under pressure. The density profiles in panel **e** showcase a situation where Pucc is kept at its equilibrium value 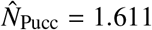 of panel **d**, see Supplementary Fig. 4, while 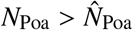 and 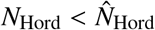. We observe zonation, discontinuities, and competitive exclusion along the salinity gradient.

**Supplementary Fig. 4.**
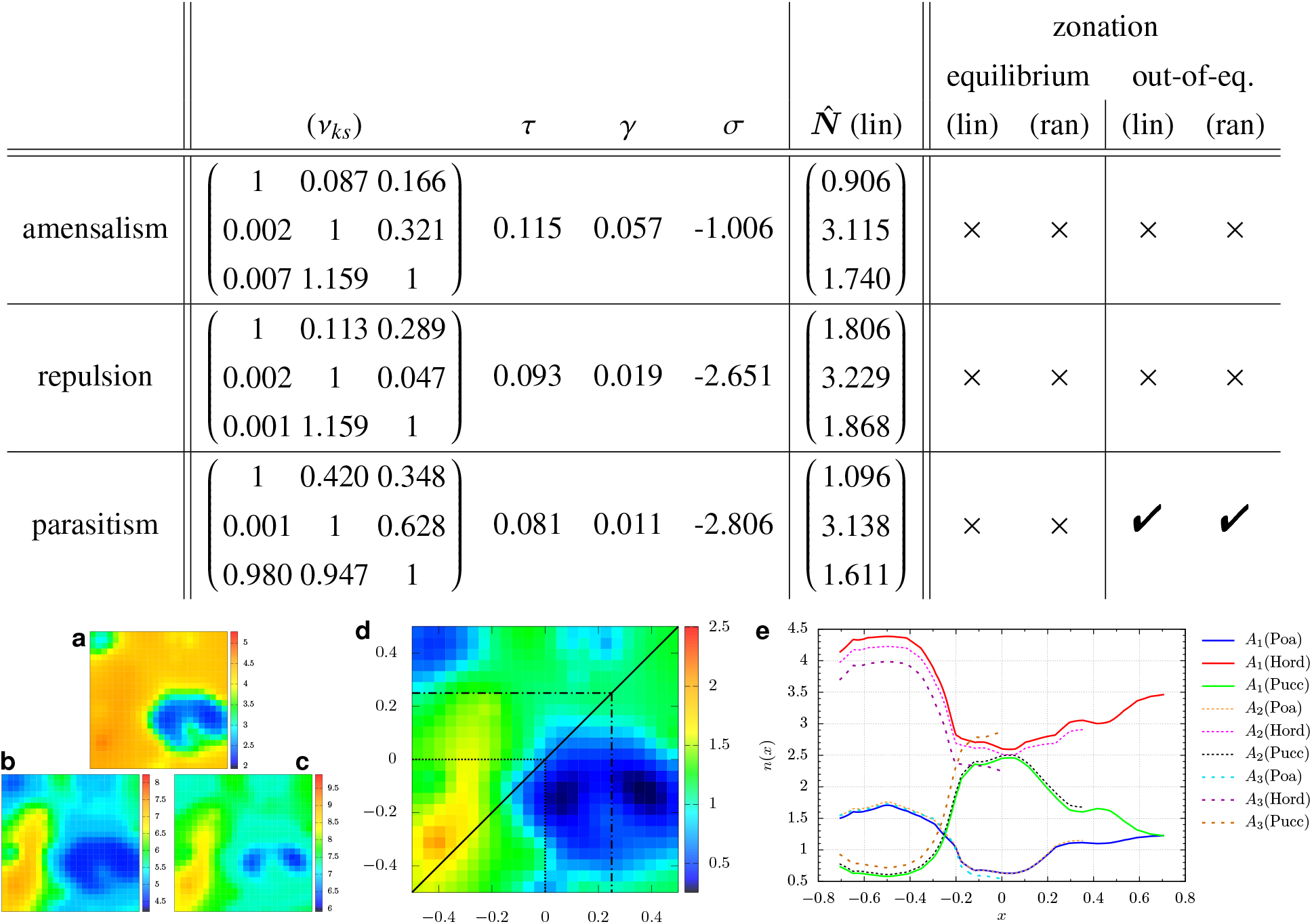
Fitted parameters and further details on the DFTe model. Fitting the energy function of equation (29) to the mixture abundances shown in Supplementary Fig. 3a, we obtain the six off-diagonal resources requirements *ν_ks_*, the dispersal coefficient *τ*, the interaction strength *γ*, and *σ* for the resource weights. The three resources associated with a randomly generated salinity landscape are depicted in panels **a**–**c**. The equilibrium density distribution of Poa, depicted in panel **d**, deviates from its limiting resource *R*_1_ (**a**) due to two mechanisms: (i) the competition with Hord and Pucc, and (ii) the intra-specific dispersal pressure that prohibits complete utilisation of the provided resources. We identify the quantitative impact of either of the two mechanisms by comparing panel **d** here with panel **a** of Extended Data Fig. 2, where Poa in isolation maximises the consumption of its limiting resource and is only restrained by the intra-specific dispersal pressure. Panel **e** shows the densities along a cut (solid diagonal line of panel **d**) through area *A*_1_ = *A* for all three species and reveals some quantitative (albeit no qualitative) changes when equilibrating in the restricted smaller parcels *A*_2_ (*A*_3_) enclosed by the dash-dotted (dashed) lines of panel **d**. The disparities between the densities for different areas reflect the altered trade-offs between the overall resource consumption and the other contributions to *E*, while the congruence between the density patterns reflects the local nature of the interactions and resource dependencies.

#### Notes on Fig. 5

In Refs. [38, 39], Veilleux reports the abundances of *Paramecium aurelia* (P), which feeds exclusively on Cerophyl, and *Didinium nasutum* (D), which feeds exclusively on P, under various conditions over periods of weeks. Supplementary Fig. 5 illustrates the three time series (out of five datasets in Ref. [38]) that exhibit the most unambiguous cyclic trajectories. Arguably, only Supplementary Fig. 5a shows a more or less stable orbit over several periods and can thus be taken seriously as a predator–prey cycle. We deem the other two time series in Supplementary Fig. 5b/c not to qualify of having reached steady-state dynamics due to (i) the considerable disparity between the last and second-to-last complete cycle of the dataset shown Supplementary Fig. 5b, and (ii) the upward trend of P in Supplementary Fig. 5c.

**Supplementary Fig. 5.**
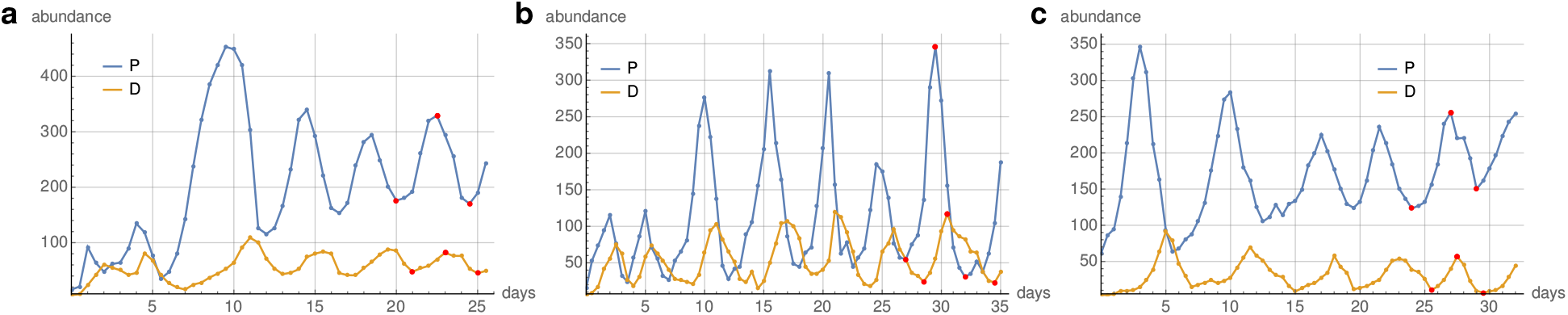
Measured predator–prey cycles. Veilleux’s experimental data of the time evolution of abundances for *Didinium nasutum* (D) feeding on *Paramecium aurelia* (P) [38, 39]. Panels **a**, **b**, and **c** reproduce Figs. 14c, 11a, and 12a of Ref. [38]. The straight line segments guide the eye. We assume the steady-state abundance oscillations to cycle around reference equilibrium abundances 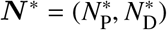, which we estimate as arithmetic means of the minima and maxima (red dots) of the last fully visible cycle of each time series, see Supplementary Tab. 1. We refrain from over-interpreting ***N***^∗^, which merely anchor our comparison between the trajectory on 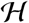 and the experimental data. The last full cycle in panel **a** presents the most convincing steady-state trajectory among all three time series and serves as our primary benchmark.

We aim at predicting the final cycle of all three datasets shown in Supplementary Fig. 5 with DFTe, while bearing in mind that any ‘steady-state’ predictions for the data of Supplementary Figs. 5b/c have to be taken with a grain of salt. With both species in suspension, we face a uniform situation like for Tilman’s experimental setup studied above. Cerophyl concentrations are given in units of 1.8 g/litre. The unit of energy is fixed by *ζ* = 1, see equation (7), and length by *A* = 1000. The Cerophyl resource for P is maintained at constant concentration *ρ_c_*, while P itself serves as resource for D. The data in Ref. [39] show that the abundance of P in monoculture scales linearly with *ρ_c_* (*N*_P_ = 500 for *ρ_c_* = 0.5), such that the Cerophyl requirement of P is *ν_cP_* = 1. The prey requirement of D also depends on *ρ_c_* with more (starved) prey individuals required at lower Cerophyl concentration. All DFTe input parameters are extracted from the data reported in Refs. [38, 39] and listed in Supplementary Tab. 1. The predator–prey relation between the two species is certainly beneficial for D and harmful for P, such that parasitism is the natural choice for the interaction functional, see Extended Data Tab. 1. That is, excluding self-interaction and installing D as the logical beneficiary of the interaction, we set up the competition matrix *γ_ss′_* = *γ δ_sD_ δ*_*s′ P*_ with only one nonzero component. Disregarding the experimental uncertainties in the resource requirements of P and D, we are left with the interaction strength *γ* as sole fit parameter in equation (30). In Supplementary Tab. 1 we report *γ* = 29.2 for Supplementary Fig. 5a to yield the best match between 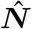 and the reference equilibrium abundances ***N***^∗^. The according DFTe hypersurface in Extended Data Fig. 3a not only produces a reasonable match between the abundances predicted by DFTe and ***N***^∗^, but also between the last cycle of Supplementary Fig. 5a and an equipotential line (red) of appropriate amplitude on 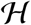. Optimising the resource requirement of D within its experimental range of uncertainty, we obtain the hypersurface in Fig. 5 and Extended Data Fig. 3b, which shows an even better match between data and DFTe calculation. The comparison with the squeezed equipotential line in Extended Data Fig. 3c suggests that, as common sense demands, parasitism outperforms amensalism in describing the predator–prey oscillations with DFTe.

**Supplementary Tab. 1.**
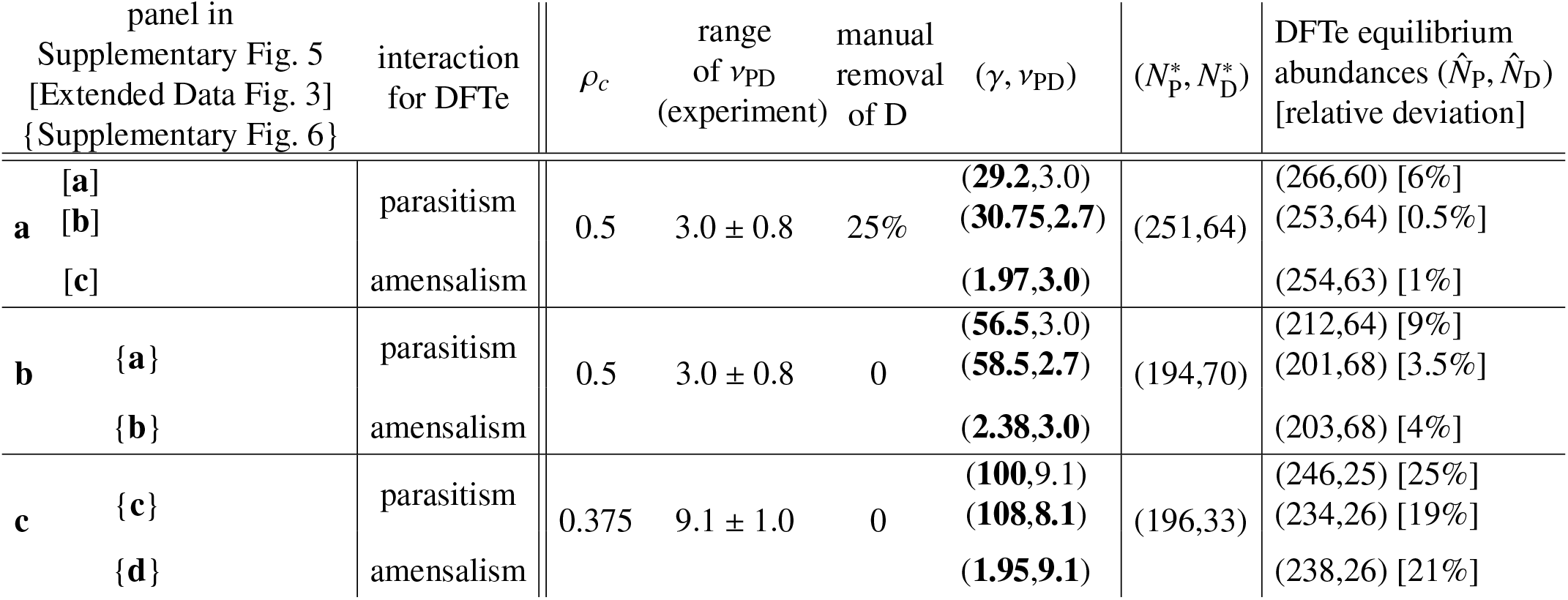
Fitted parameters and further details on the DFTe model. The minima of the DFTe hypersurface, built from equation (30) with the best fits for the two parameters *γ* and *ν*_PD_, are most consistent with Veilleux’s data for a parasitic relation between predator and prey. As figure of merit we choose the relative deviation between the DFTe equilibrium abundances and 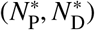, see Supplementary Fig. 5. Fitted parameters are typeset in bold. In the experiment represented by Supplementary Fig. 5a, D is manually removed from the suspension on a regular basis, corresponding to a *Δ* = 25% reduction of D, which is implemented in equation (30) by the replacement *N*_D_ → *N*_D_/(1 − *Δ*). If we refrain from optimising D’s resource requirement *ν*_PD_ within its experimental uncertainty and stick with its central value, we still find the according one-parameter predictions of DFTe to match the data-based estimates 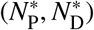 reasonably well. We tried (unsuccessfully) to improve the results for amensalism [encoded by the replacement 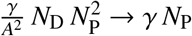 in equation (30)] by adjusting *ν*_PD_ away from its experimental central values. The DFTe equilibrium abundances correlate best with the experimental data depicted in Supplementary Fig. 5a, which incidentally represents the most convincing of the predator–prey cycles reported in Refs. [38, 39]. While the equilibrium abundances presented here do not justify making a strong case for either parasitism or amensalism, we settle the issue in favour of parasitism by observing the cycles on the DFTe hypersurface away from its minimum, see Extended Data Fig. 3 and Supplementary Fig. 6.

**Supplementary Fig. 6.**
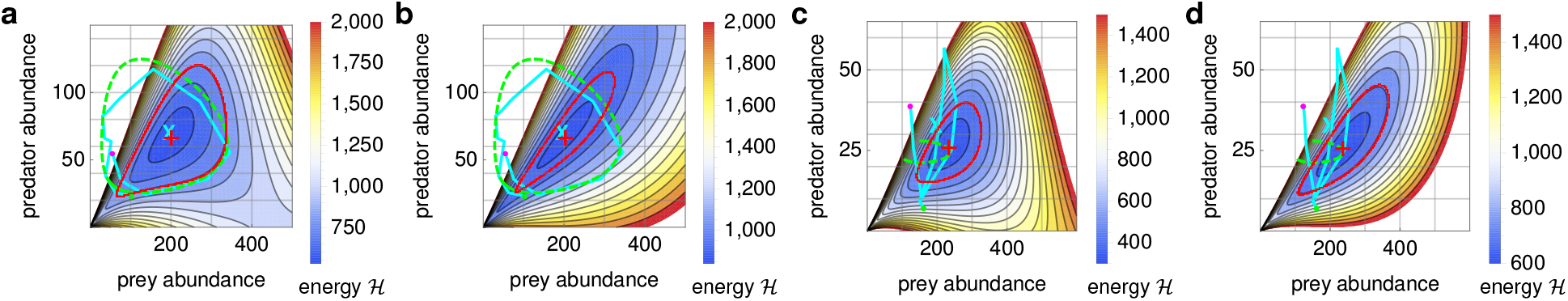
DFTe predictions for two less stable predator–prey cycles. Although the alleged steady-state cycles depicted in Supplementary Fig. 5b/c are not stabilised towards the end of the time series, we find a reasonable match with an equipotential line of the DFTe hypersurface 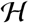 at least for panel **a**, which benchmarks DFTe against the data of Supplementary Fig. 5b with the parameters given in Supplementary Tab. 1. Panel **b** demonstrates that the same data cycle is less well captured if we choose amensalism instead of parasitism. The analogous calculations for Supplementary Fig. 5c are illustrated in panels **c** and **d**. Comparing panels **a** and **c**, we observe that a disturbance of the cycle can lead to the extinction of D more easily at higher Cerophyl concentration *ρ_c_*. This is in line with the experiments reported in [39], where no stable cycles are found for *ρ_c_* > 0.75. In view of the upward trend of P in Supplementary Fig. 5c near the end of the time series, we may argue that the equilibrated 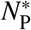 is somewhat higher than what the cyan cross indicates, which would bring the DFTe equilibrium abundances in better agreement with the data. The Lotka–Volterra model (dashed green lines), see Supplementary equation (S17), performs well in panels **a**/**b** and fails in panels **c**/**d**. However, we do not want to over-interpret these observations, bearing in mind the experimental uncertainties pertinent to the steady-state character of the two setups of Supplementary Fig. 5b/c.

The equipotential lines drawn in Extended Data Fig. 3 and Supplementary Fig. 6 are meant to test the predictive power of the DFTe hypersurface away from its minimum. A quantitative time-dependent version of DFTe, in particular the functional form of the resource energy (which is simply quadratic in equation 9), has to be informed by dynamic variables and characteristic time scales such as birth- and death rates, time delays in interaction mechanisms (for example, due to metabolism or foraging), environmental variability and so forth. The predator–prey cycle can approximately be regarded as an ellipse, which is determined by four parameters (position, orientation, and eccentricity). Our use of three fit parameters (*ν*_PD_,*γ*, and *ΔE*) would thus not suggest a particularly powerful approach. However, the underlying functional in equation (30) provides causal understanding of the intra- and inter-specific relations and follows from the universal DFTe functional in equation (9), which we proved to be applicable to a wide range of systems — the significance of our DFTe approach stems from its mechanistic and unifying qualities. Furthermore, the three-parameter DFTe equipotential lines in Fig. 5, Extended Data Fig. 3, and Supplementary Fig. 6 are of a quality similar to the Lotka–Volterra predictions from

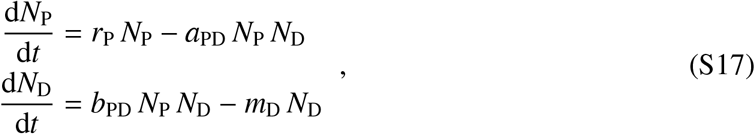

where *six* parameters yield best fits to the experimental cycles of *N*_P_*, N*_D_ in Supplementary Fig. 5 for

| Supplementary Fig. | $N_P(0)$ | $N_D(0)$ | $a_{PD}$ | $b_{PD}$ | $m_D$ | $r_P$ |
| --- | --- | --- | --- | --- | --- | --- |
| 5a | 171.119 | 61.5892 | 0.01962 | 0.005960 | 1.41462 | 1.25682 |
| 5b | 46.1652 | 34.5678 | 0.02552 | 0.005417 | 0.67998 | 1.59854 |
| 5c | 108.445 | 22.0646 | 0.09895 | 0.001345 | 0.20560 | 2.41680 |

Here, *N*_P_(0) and *N*_D_(0) are the fitted initial prey and predator abundances, respectively, *t* is time, *r*_P_ is the intrinsic growth rate of the prey, *a*_PD_ is the per-predator per-prey predation rate, *b*_PD_ is equal to *a*_PD_ times a conversion factor, and *m*_D_ is the mortality rate of predators.

#### Notes on Fig. 6

In the following, we provide a detailed account of the DFTe input and the resulting density distributions shown in Extended Data Fig. 6. The according simulations, which eventually lead to Fig. 6, always include dispersal (*E*_dis_[***n***]), whose trade-off against resources (*E*_Res_[***n***]) creates an intuitively accessible base line for studying more complex (sub-)systems that include environments (*E*_env_[***n***]) and interactions (*E_γ_*[***n***]).

Comparing the Tree distribution *n*_5_ in Extended Data Fig. 6a (‘**a**’ in the following) with the density in Extended Data Fig. 5e, we observe the effect of the inter-species competition over resources. Without any further interactions, all three species {1(Fungus), 5(Tree), 6(Grass)} have the same ability to exploit the resources (since *τ_s_* = 0.01, see Extended Data Tab. 3), resulting in *n*_5_ = *n*_6_ (both species have the same resource requirements). The Fungus does not require sunlight (*ρ*_5_) and finds its primary niche near the western boundary, where (i) water and nutrient occur at high density and (ii) the Tree and Grass exert little competition pressure due to their small densities induced by a lack of sunlight. Although the Tree and Grass are identically parameterised in **a** (prior to the inclusion of environments and interactions), they are still distinguishable species because the (***n***-)nonlinearity of equation (31) implies different energies for (i) two species with independent variables *n*_5_ and *n*_6_ versus (ii) one ‘effective species’ comprised of the sum *n*_5_ + *n*_6_. All three species have the same density profiles in the south-eastern quadrant, where water is their limiting resource. Both Tree and Grass are limited by sunlight in the west and thus must let the Fungus take advantage of the underutilised resources of water and nutrient, cf. *ρ*_lim_ in Extended Data Fig. 5. For the central ring region we observe that the Tree and Grass equally divide their limiting resource (sunlight, at densities of ~ 9 … 15) and therefore aim at extracting an equivalent amount of nutrient. This puts competition pressure on the nutrient-limited Fungus, resulting in *n*_1_ ≈ 4 < *n*_5_ = *n*_6_ ≈ 6, despite Tree and Grass being constrained by three resources while the Fungus only requires two.

As the environments are added in **b**, the Fungus distribution (and, consequently, its abundance *N*_1_) changes only marginally since its density in the environmentally hostile regions is low anyway. In contrast, 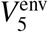 expels the Tree from the hostile central region, allowing the Grass to invade. This includes the central ring region, where the Grass environment 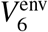 is also hostile. There, the Grass density even exceeds that of the neutral (uniformly zero) environment employed in **a**. This phenomenon of an increased density in a less favourable habitat originates in the trade-off between all DFTe energy components of all species and may be perceived as counterintuitive prior to our quantitative assessment.

The picture of a Grass-dominated landscape changes drastically as we complete the subsystem of the species {1, 5, 6} by introducing their bipartite interactions in **c**. The combined resource-environment-dispersal constraints on densities are modified by the mutualism between Tree and Fungus, which permits higher densities of both species and tends to align them. As a result, Grass comes under considerable pressure due to (i) the amensalism where Trees exist and (ii) the repelling Fungus. While the Fungus-Grass repulsion affects both species in a symmetric way (and allows Grass to expel the Fungus from its low-density regions), the Tree-Fungus alliance renders Grass competitively inferior to the Fungus in regions of high Tree density. Grass can coexist with both the Tree and the Grass (where their densities are low) and even dominates in regions that are both poor in resources and environmentally hostile for the Tree. However, we predict competitive exclusion of Grass from Fungus territory, except where (prior to interactions) the Fungus density is very low and the Grass density is very high. This balance between species, enforced at the DFTe energy minimum, also gives rise to sharp interfaces between Grass and the other two species, contrasting with the smooth (noninteracting) densities in **a** and **b**. Note that this zonation pattern is equilibrated, unlike the transient zonation states in Extended Data Fig. 2. The zonation originates in the Fungus-Grass repulsion but is transferred to the Tree distribution via the Tree-Fungus mutualism. More of such discontinuities emerge and transform as we keep adding species and their interactions.

In **d** we introduce the Deer, who owes its relatively uniform distribution to the lack of adversarial interactions like those between Grass and Fungus. Although all heterospecifics suffer from the additional competitor over resources, the Tree nearly makes up for the loss through the added mutualistic interaction. That is, the Deer’s resource consumption primarily lowers the densities of the Fungus and (to a lesser extent) of Grass, which alleviates both species’ conspecific dispersal pressure and the Fungus-induced repulsive pressure on Grass. The result is a less pronounced zonation between Grass and its competitors. Although the Grass density decreases, its land cover increases where the Fungus recedes. In summary, the global Tree and Grass abundances are roughly maintained, and the main impact of introducing the Deer is the (a priori surprising) decline of the trophically distant Fungus.

The Snail, introduced in **e**, has the same habitat preference and resource requirements as the Fungus. As expected, the resulting intense competition between the two species is decided in favour of the superior Snail, see Extended Data Tab. 3, which expels the Fungus from favourable regions. However, the mutualistic trio with the Tree and Deer protects the Fungus against the Snail in areas of high Tree density. While the competition via asymmetric interaction also incurs a cost for the Snail, albeit lower than for the Fungus, the Snail more than compensates this disadvantage (compared to the mutualistically bolstered Tree and Deer) through its low dispersal pressure, which enables a more efficient exploitation of resources. This benefits the Snail particularly in the heavily contested central ring region, where a high Grass density (implied by the excluded Fungus and diminished Tree) supports the Snail commensalistically. Both Deer and Tree, mutualistically connected to the Fungus, suffer from lowered Fungus density. Aside from occupying former Fungus territory, the Snail can therefore carve out the two ‘circle segments’ beside the central ring region, where all the other species are relatively low in density (before and even more so after the Snail’s introduction) and thus offer little resistance. Although Grass does not benefit from the Snail through a direct interaction, the implications of the Snail’s introduction ripple through the community and make Grass the main beneficiary, particularly where the Fungus and the Tree recede along with their competitive pressure.

The Pig, introduced as predator in **f**, relies on and thus aligns with the Fungus. The cost (*E*_Res_) of nonzero Pig density in regions void of Fungus can only be absorbed to a minor degree by the other energy components (in particular the nonlimiting resources), such that the Pig is largely restricted to Fungus habitat. The predation has several implications across the community. As expected, the Fungus density is severely diminished globally, and more of the resource pools *ρ*_4_ and *ρ*_6_ in the affected regions become available to the rest of the community. The Pig consumes only a small part of this excess *Δρ*_4,6_, and both the Snail and the Grass do not benefit from the released resources since their habitat hardly overlaps with that of the Fungus. The Tree loses most of its mutualistic Fungus support, most visibly in regions of high Tree density, which outweighs the positive impact of *Δρ*_4,6_. The alleviated amensalistic pressure from the Tree not only allows Grass to invade into former Tree territory (aided by the diminished repulsion from the Fungus), but also releases additional resources, incidentally in some of the preferred habitat of the Deer and Snail. The combination of these direct and indirect mechanisms thus leaves the Deer, the Grass, and especially the Snail as the beneficiaries of the Pig’s introduction. While the positive impact of the Pig on Grass and Snail are in line with an intuitive examination of the interactions illustrated in Fig. 6, the growth in the Deer population, mutualistically connected to the declining Tree, may come as a surprise if not all the ecosystem ingredients are taken into account quantitatively.

In **g**, we finally introduce the Cat as the apex predator, with an equilibrated density distribution that is maintained by five Deer and one Pig per individual, see Extended Data Tab. 3. The Cat closely aligns with the Pig, except where the Deer is the limiting prey that alleviates the predation pressure on the Pig (such as the eastern circle segment, where the ratio between Deer and Pig falls below 5:1) by making the Pig a nonlimiting resource that is affected only marginally due to *σ* = −4 for the resource weights. The Fungus distribution correlates with the Pig population not because of a causal relationship — if anything, we would expect the Fungus to benefit as the Pig becomes prey for the Cat. The Pig distribution follows the Fungus distribution as per their predator–prey relationship, and the Fungus is in decline because (i) the Tree is receding and (ii) the Snail establishes a stronger foothold globally, also in the western Fungus habitat. The source of this cascade of changes in the community is the heavy predation of the Deer, accompanied by a decline of the mutualistically connected Tree, which therefore decreases its amensalistic pressure on Grass and its support for the Fungus. The Snail then benefits from an elevated Grass density as well as the released resources and puts additional competition pressure on the Fungus, such that the Tree loses even more support. This negative (positive) feedback loop between Tree, Deer, and Fungus (Grass and Snail) finally stabilises at the DFTe energy minimum with the density distributions in **g**. By exterminating the Cat (that is, transiting from **g** to **f**), we affect the community the opposite way — the time-independent DFTe energy functional in equation (31) does not include history and therefore no hysteresis. In Fig. 6 we depict the relative changes 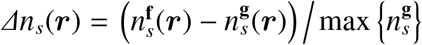 of the equilibrium density distributions for the species *s* ∈ 1–6.

If we were to strengthen the Cat population, we could, for example, reduce deforestation globally. This benefits (i) the Deer and (ii) the Pig (through enhancing the primary Fungus habitat, see first panel of Extended Data Fig. 7a). The quantitative impact of

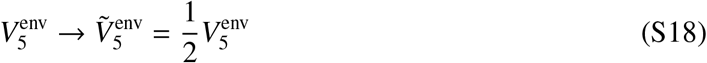

is shown in Extended Data Fig. 7b/c, and the relative density differences to column **g** of Extended Data Fig. 6 are shown in Fig. 6c.

Approximating realistic long-range interactions by contact-type (i.e., zero-range) interactions, as we implicitly do in our synthetic food web example, is only justified if the simulation area *A* is sufficiently coarse-grained into areas *A*′ < *A*, such that regions beyond the focal area *A*′ (which contains the focal position ***r***′) have negligible interaction effect on the density *n*(***r***′):

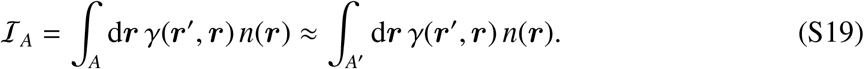

This requirement crucially depends on the interaction kernel *γ*, whose characteristic length scales relate to *A*. As an illustration, we take a Yukawa potential *γ*(***r***′, ***r***) = *γ* exp[−|***r*** − ***r***′|*/l*]/|***r*** − ***r***′| with characteristic length scale 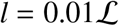, focal position ***r***′ = 0, and constant densities *n*(***r***) = *n* in a disk-shaped area 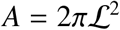, such that 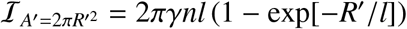. Then, demanding 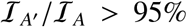, i.e., obeying Supplementary equation (S19) accordingly, requires us to choose 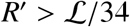. Note that *A* in Fig. 6 is subdivided into parcels of 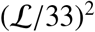.

#### Outlook: DFTe-based dynamics across temporal scales

Aiming at the simultaneous integration of ecosystem drivers on all time scales, we propose to extract the ecosystem dynamics from trajectories in the space of DFTe energies. Inspired by the mathematical structure of DFTe, we identify two promising routes for addressing time evolution on the (short) dispersal and the (intermediate) ecological time scale, respectively: We propose that processes on the dispersal time scale enter our formalism at the level of the selfconsistent equations of DPFT, specifically through an alignment energy (akin to dipolar interactions of quantum gases) that iteratively steer the directed flow of nonequilibrium density distributions into the states of lowest energy for fixed abundances. Encouraged by our results on Veilleux’s predator–prey system in Fig. 5, we also hypothesise that the set 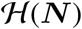 of these states, spanned by all possible abundances ***N***, is the platform on which time-dependent processes on the ecological time scale evolve through Newtonian-type dynamics. The simultaneous temporal drivers of ecosystem dynamics are drift (stochasticity of densities and abundances), speciation (emergence of new species), environmental changes, and time-dependent resources — in addition to temporal properties of species, such as situation-specific dispersal through space, growth rates, and time lags in response to stimuli [4, 44]. These drivers exert their influence on different time scales, which overlap depending on the involved processes and species. For example, speciation accompanies the evolutionary time scale, which is often tagged to long-term geological processes and climate. Drift happens at shorter time scales, such as those associated with the rate at which individuals disperse through space, but its accumulated effects can influence long-term features like extinction rates. The intermediate ecological time scale is characterised by population changes induced by reproductive cycles, growth and predation rates, and accompanies short-term environmental variability such as seasons, weather, cycles of day and night, or tides. See Ref. [53] for a review on transient phenomena in ecology that stretch across these time scales.

We propose to capture this overlapping trinity of time scales by simultaneously augmenting the DFTe framework with three types of time evolution. The according original hypotheses and mechanisms proposed below are a road map for creating a universal tool that predicts the nonequilibrium dynamics of complex ecosystems across all scales of time and space.

To establish a universally suitable starting point for ecosystem dynamics, we propose to identify datasets that suggest adiabatic evolution (a succession of time-independent equilibria for slowly changing external parameters), which drags the system along with the evolving equilibrium. We can then model the ‘external dynamics’ of such time-(*t*-)dependent environments ***V***^env^(*t*) and resources ***ρ***(*t*) as extensions of their static versions by fitting and predicting density data in the fashion of Figs. 2–6. For example, the measured response of a species *s* to a climatic change yields a parameterised 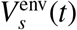 for modelling natural settings — much like the quadratic environment in equation (24) is informed by the species’ static response to a temperature gradient in a simple setup and used in a complex setup. These *t*-dependencies (potentially, on all time scales) promote the global minimum 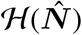 of *E* to a time-dependent attractor 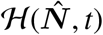. First targets should be systems with negligible speciation and drift. Both mechanisms can eventually enter the time-dependent space of DFTe energies *E*[***n***, ***μ***](***N***, *t*) (of which 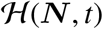 is one slice and 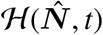 one point) via explicit *t*-dependent recipes at the level of external dynamics and/or the DPFT loop of equation (21). Note that speciation and extinction manifest in a time-dependent dimensionality of 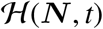. This agenda will either render the existing functional structure of *E* sufficient or identify refined forms of its components.

The time dependence of densities and abundances arise through external dynamics on all time scales, superimposed with species-specific ‘internal dynamics’ on the dispersal and ecological time scale. We propose to capture the internal dynamics with two internal time-evolution mechanisms. One is induced by the selfconsistent DPFT loop of equation (21) itself, which iteratively transforms nonequilibrium density distributions ***n*** into the state of lowest energy for given abundances ***N***. This form of time evolution implicitly contains a dissipative mechanism that permits the existence of a trajectory towards the DFTe hypersurface 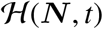. Suppose, for example, that a drift event removes one out of *N* individuals. The formerly equilibrated density distribution is now out of equilibrium and will approach the new attractor 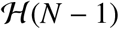 with a rate that is determined by time-dependent species-specific admixing parameters *θ_s_*(*t*), see equation (21). Their *t*-dependence encodes how well each species copes with an altered (effective) environment and has to be informed by real data. We mean this ‘*θ*-equilibration’ to exert its influence primarily on the dispersal time scale, where a fixed abundance is redistributed due to (and in proportion to the magnitude of) short-term disturbances. This includes the scenario of a perpetually changing 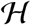, for example due to periodic driving, which can prohibit the system from ever *θ*-equilibrating. Inspired by the formalism of fluid dynamics, we may establish a velocity field ***v***(***r***, ***θ***) = {*v*_1_(***r***, *θ*_1_(*t*)), …, *v_S_*(***r***, *θ_S_* (*t*))} that connects successive density distributions. Such a velocity field describes the dissipative flow of the density distribution towards its lowest-energy configuration on 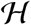 and can itself put constraints on the DFTe energy functional: We hypothesise that this back reaction supplements the (either static or explicitly *t*-dependent) DFTe energy with a velocity-dependent component *E_**v**_*, which vanishes at *θ*-equilibrium, where |***v***| = 0 everywhere, but which favours specific velocity alignments during *θ*-equilibration. Such a dynamic energy component selfconsistently steers the flow of densities by balancing the cost of ‘misaligned’ velocity vectors (inspired, for example, by dipole-dipole interacting quantum gases) against all other energy components. This agenda aims at predicting population redistributions on the dispersal time scale, such as the flow of migrating wildebeests, swarming locusts, or schools of fish, see Supplementary Fig. 7. Related velocity fields of microswimmers in a hydrodynamic setting have been studied [54].

**Supplementary Fig. 7.**
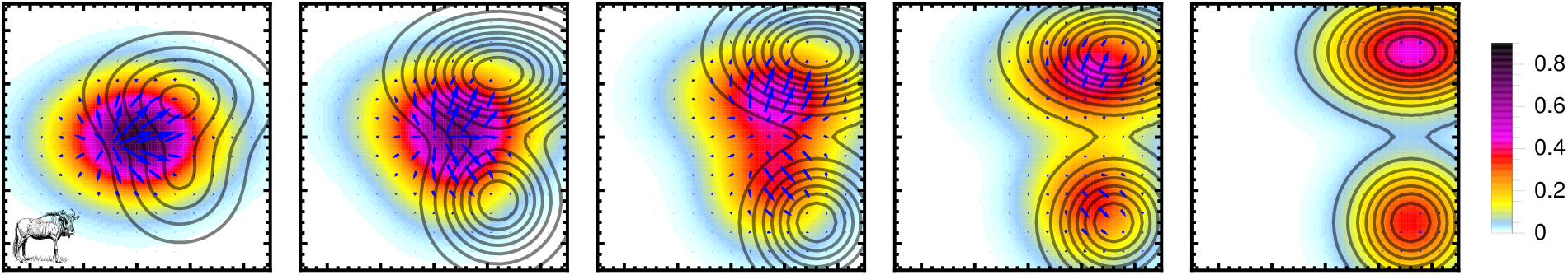
The five panels illustrate (snap1shots of) the velocity field ***v*** (blue arrows), which (i) is associated with the flow of a hypothetical density distribution of wildebeests (colour-coded), (ii) is induced by a time-dependent environment (precipitation, contour lines), and (iii) accompanies the *θ*-equilibration of the density distribution (towards the right panel).

We propose to supplement the explicit external dynamics and the *θ*-equilibration with a third mechanism of time evolution that operates predominantly on the ecological time scale. Motivated by our predator–prey results presented in Fig. 5, we argue that species-specific growth rates constrain the possible trajectories that a system can take on 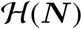. In particular, a steepest descent along 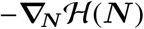 towards 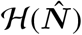 or even the approach of 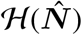 along any trajectory may be forbidden. The latter scenario entails an ‘excitation’ energy *ΔE* above 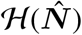, which cannot be dissipated. Then, *ΔE* implies steady-state cycles and terminal trajectories in predator–prey systems akin to those hinted at in Fig. 5, or oscillating abundances due to time lags between resource consumption and growth. In Fig. 5, we observe abundance changes over intermediate time scales due to birth and death in an environment that changes slowly enough for 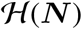 to be considered static. Under the assumption that the dispersal time scale is much shorter than the ecological time scale, the density distributions *θ*-equilibrate to 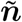 before significant changes in ***N*** occur, such that we trace a trajectory on 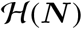. Of course, in reality the events and mechanisms of the different time scales are at work simultaneously. We may single out the ecological time scale only if it is well separated from the evolutionary time scale and if short-term variations at the dispersal time scale are negligible. Indeed, these conditions are met by Veilleux’s predator–prey steady states, which evolve in a uniform time-independent environment void of evolutionary events.

We suggest to extract the relationship between generational rates (e.g., birth or predation rates) and *ΔE* by matching time-series data with suitable equipotential lines 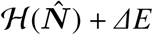 on a static 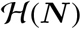. Generally, *t*-dependent excitations *ΔE*(*t*) accompany a *t*-dependent 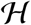 that is, for instance, driven by *t*-dependent environments. Analogous to particles in Newtonian mechanics, we can propagate the abundances ***N*** through the recipe

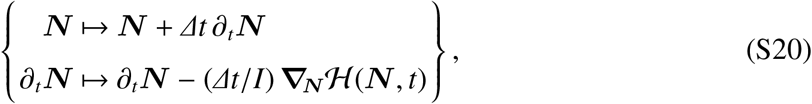

which allows us to extrapolate limited time series data. Given an energy functional *E*[***n***, ***μ***](***N***, *t*), the sole fit parameter in the ‘***N*** -evolution’ of equation (S20) is *I*, which encodes an inertia of the system akin to mass. Discrepancies between ***N*** -evolution and experimental data will help to refine *E* and, consequentially, the driving forces induced by the slope of 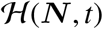, towards a universal energy functional that endows the geometry of 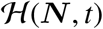 with predictive power for time evolution on the ecological time scale.

#### Image sources

We obtained permission to display the following photographs of Fig. 1: Fig. 1c. *Fragilaria crotonensis* (top left), *Asterionella formosa* (bottom left), *Tabellaria flocculosa* (right), copyright owner: Jason Oyadomari. *Synedra filiformis* (centre), copyright owner: Don Charles.

Fig. 1d. *Poa pratensis* (left), copyright owner: Igor Sheremetyev. *Puccinellia nuttalliana* (right), copyright owner: Katy Chayka.

The silhouettes in Fig. 1 and Supplementary Fig. 7 come with public domain licence:

- https://publicdomainvectors.org/en/free-clipart/Red-mushroom-with-dots/76883.html
- https://publicdomainvectors.org/en/free-clipart/Vector-clip-art-of-bear-from-the-Flag-of-California/17297.html
- https://publicdomainvectors.org/en/free-clipart/Tree-silhouette-vector-graphics/7129.html
- https://publicdomainvectors.org/en/free-clipart/Buck/42510.html
- https://publicdomainvectors.org/en/free-clipart/Wild-boar-silhouette/55391.html
- https://publicdomainvectors.org/en/free-clipart/Vector-illustration-of-snail/30685.html
- https://publicdomainvectors.org/en/free-clipart/Leopard-vector-graphics/35615.html
- https://www.kisscc0.com/clipart/wildebeest-drawing-line-art-cartoon-public-domain-umvghc/

